# Light-activated tetanus neurotoxin for conditional proteolysis and inducible synaptic inhibition *in vivo*

**DOI:** 10.1101/2025.01.27.635161

**Authors:** Heegwang Roh, Dongwook Kim, Byeongchan Kim, Younghyeon Jeon, Yeonghye Kim, Martin Jacko, Fei Xu, Chang Lin, Ji Won Um, Alice Y. Ting

**Author notes:** Howard Hughes Medical Institute, University of California at Berkeley, Berkeley, CA 94720, USA. Aperture Therapeutics, 733 Industrial Road, San Carlos, CA 94070, USA. Department of Genetics, Yale School of Medicine, New Haven, CT 06510, USA. H. Roh and D. Kim contributed equally to this work. Correspondence (A.Y.T.), (J.W.U.).

## Abstract

The light chain of tetanus neurotoxin (TeNT) is a 52 kD metalloprotease that potently inhibits synaptic transmission by cleaving the endogenous vesicle fusion protein VAMP2. To mitigate the toxicity of TeNT and harness it as a conditional tool for neuroscience, we engineered Light-Activated TeNT (LATeNT) via insertion of the light-sensitive LOV domain into an allosteric site. LATeNT was optimized by directed evolution and shown to have undetectable activity in the dark mammalian brain. Following 30 seconds of weak blue light exposure, however, LATeNT potently inhibited synaptic transmission in multiple brain regions. The effect could be reversed over 24 hours. We used LATeNT to discover an interneuron population in hippocampus that controls anxiety-like behaviors in mouse, and to control the secretion of endogenous insulin from pancreatic beta cells. Synthetic circuits incorporating LATeNT converted drug, Ca^2+^, or receptor activation into transgene expression or reporter protein secretion. Due to its large dynamic range, rapid kinetics, and highly specific mechanism of action, LATeNT should be a robust tool for conditional proteolysis and spatiotemporal control of synaptic transmission *in vivo*.

## Introduction

Neurons and multiple other cell types secrete signaling molecules into the extracellular space to mediate intercellular communication and maintain homeostasis^1, 2^. These cells use a conserved mechanism of secretion involving SNARE complexes that link proteins such as VAMP2 on the surface of exocytic vesicles with SNAP25 on the plasma membrane^1^. SNARE complexes bring the two membranes into close apposition and drive membrane fusion. Due to the central and conserved role of VAMP2 and SNAP25 in mammalian biology, bacterial toxins have evolved highly specific mechanisms to interfere with these two proteins and prevent secretion, thereby poisoning the mammalian host. For example, tetanus neurotoxin from the bacterium *Clostridium tetani* targets VAMP2 via a metalloprotease encoded in its light chain that is delivered into the cytosol of neurons via a receptor-binding and translocation-competent heavy chain^3^. This design makes TeNT extremely potent; as little as 0.2 ng/kg is lethal to humans^4^.

Despite TeNT’s extreme neurotoxicity, its highly specific mechanism^5–8^ which produces cleavage of only a single protein (VAMP2 and its isoforms) in the mammalian proteome^9^ has made TeNT a powerful platform for tool development. The protease light chain of TeNT (52 kD) has been used to silence synaptic transmission in specific cell populations, leading to discovery of functional circuits in spatial learning and social behaviors^10, 11^. Additionally, TeNT and the related toxin BoNT (botulinum neurotoxin, from *Clostridium botulinum*) have been harnessed to deliver protein cargoes across the plasma membrane in a cell-type specific manner^12–16^, and BoNT has been engineered to recognize and cleave non-SNARE intracellular proteins such as PTEN^17^.

To fully realize the potential of these toxins as tools for biology and neuroscience, however, it is necessary to introduce levers for both spatial and temporal control over their protease activities. A light-regulated TeNT that could be turned on within seconds in specific regions of the brain would enable functional circuit dissection with high precision and minimal toxicity. Such a construct could also enable exogenous control of secretion processes in other cell types, such as pancreatic beta cells that produce insulin^18^.

Here we use structure-guided protein engineering and directed evolution to produce a compact (69 kD), single-chain light-activated TeNT (LATeNT). We demonstrate LATeNT’s ability to rapidly and reversibly control synaptic transmission in the mouse brain, leading to the discovery of a hippocampal interneuron population that controls anxiety-like behaviors. Additionally, we show that LATeNT can modulate secretion of endogenous insulin from pancreatic beta cells. Finally, we leverage LATeNT’s high sequence specificity to build synthetic circuits – Ca^2+^, drug, or receptor-controlled gene transcription and reporter secretion – in HEK293T cells that lack endogenous VAMP2 expression. Due to its simple design and robust performance, we expect LATeNT to have broad utility in neuroscience and synthetic biology.

## Results

### Design and directed evolution of LATeNT

To engineer LATeNT, we considered various strategies for incorporating light-responsive domains^19^ into the 52 kD light chain protease component of TeNT. We rejected the idea of splitting TeNT protease (hereafter referred to as TeNT) and fusing the fragments to light-controlled heterodimers such as CRY2/CIBN, because reconstituted split enzymes are usually much less active than the parental full-length enzymes^20, 21^, and a two-component tool exhibits much more variability than a one-component tool^22^. We opted instead for a single-chain solution (**Fig. 1A**), similar to the light-regulated proximity labeling enzyme LOV-Turbo that we recently reported^23^. We envisioned inserting the light-sensitive LOV domain^24^ into a surface-exposed loop of TeNT to “clamp” the loop in the dark state. If the loop is allosterically coupled to TeNT’s active site, then clamping may also distort the active site and render the enzyme inactive. Blue light exposure causes LOV domain to release its C-terminal Jα helix^25^; this “unclamping” of TeNT’s surface loop may rapidly restore the active site to a native-like conformation. If successful, a LOV-containing TeNT would be a single 69 kD protein, monomeric, reversible, easily packaged in an AAV vector, and show little variation in performance across a range of expression levels.

**Figure 1.**
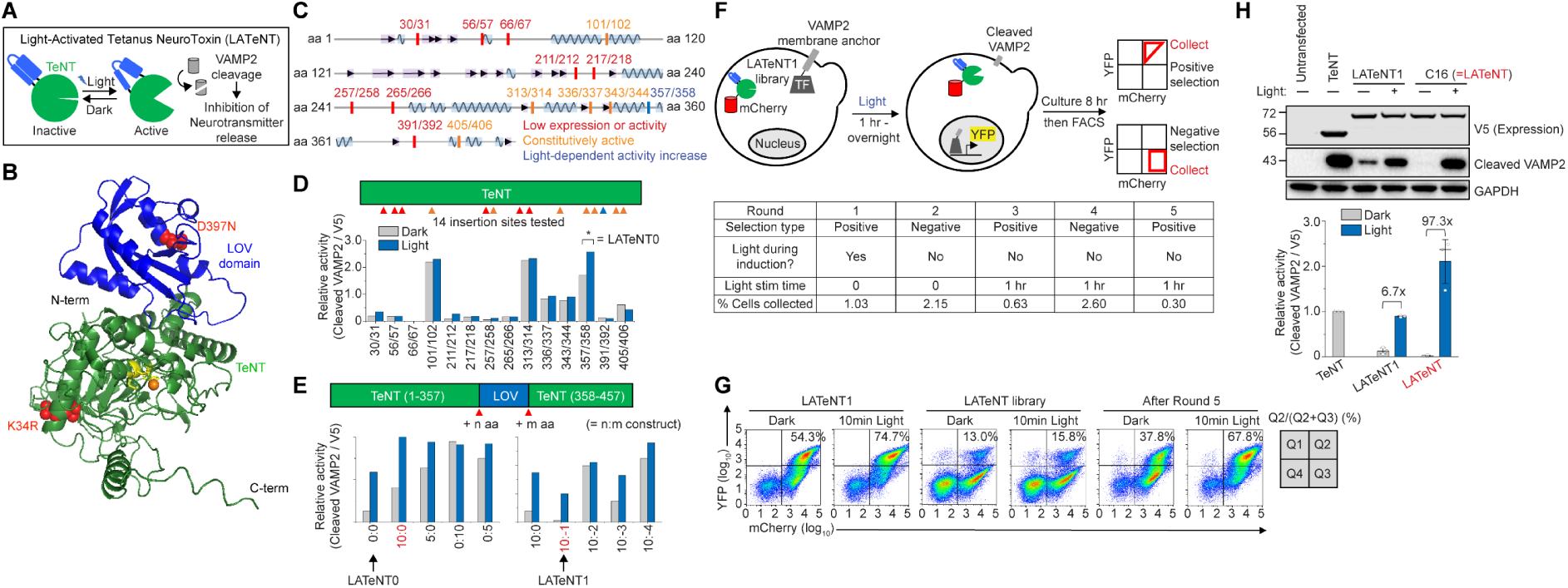
Design and directed evolution of LATeNT. **(A)** Design of LATeNT. hLOV, an engineered light-sensitive LOV domain^23^, is inserted between amino acids 357 and 358 of TeNT protease and maintains the protease in an inactive state. Protease activity is restored upon blue light (470 nm) illumination. Active LATeNT inhibits neurotransmitter release by cleaving the endogenous synaptic vesicle fusion protein VAMP2. **(B)** AlphaFold3-predicted structure of LATeNT. TeNT is shown in green, LOV domain in blue, active site residues in yellow, zinc ion in orange, and two mutations introduced by directed evolution in red. **(C)** hLOV insertion sites tested in TeNT. Color indicates result of activity test in (D). **(D)** Relative activities of LOV insertion constructs from (B). HEK293T cells expressing VAMP2 were exposed to ambient room light for 30 min. VAMP2 cleavage was quantified by Western blot. LATeNT0 was best. **(E)** Optimization of linkers in LATeNT0 produces LATeNT1. n:m indicates linkers with n and m residues N-terminal and C-terminal to the inserted LOV domain. -1 to -4 means that LOV was truncated on its C-terminal side by 1 to 4 residues. LATeNT1 was best. **(F)** Directed evolution of LATeNT1 in yeast. Cells were stimulated with light (1 hr or overnight) or kept in the dark, then sorted by YFP/mCherry ratio. Table shows conditions for each round of selection. **(G)** FACS analysis of template (LATeNT1), initial library, and post-round 5 library. Percentages indicate fraction of cells in Q2. **(H)** Comparison of LATeNT1 and final optimized LATeNT (Clone 16) in HEK293T cells after 30 min of 470 nm irradiation. This experiment was repeated 3 times. Error bars, s.d.

To engineer LATeNT in this manner, we first selected 14 sites in 12 surface-exposed loops of TeNT, based on the 2.3 angstrom crystal structure^26^, and inserted into these positions the hLOV domain, which we previously engineered for tight dark-state caging and high +light signal^27^. LATeNT candidates were expressed in HEK293T cells along with a recombinant VAMP2 reporter. Out of 14 sites tested, one site (357/358) gave a ∼1.5-fold increase in VAMP2 cleavage after 30 min exposure to room light compared to cells maintained in the dark (**Fig. 1C-D** and **Fig. S1A**). We designated this construct LATeNT0. To increase light/dark signal ratio, we then performed a linker screen to lengthen or truncate residues on either side of the inserted hLOV domain. We found that one amino acid truncation of the C-terminal Jα helix significantly improved dynamic range, perhaps by improving allosteric coupling between LOV and TeNT’s active site (**Fig. 1E** and **Fig. S1B**). Our optimized “LATeNT1” had an improved light/dark signal ratio of 6.7 (**Fig. 1H**).

To further reduce the dark state background of LATeNT1 (**Fig. 1H**) – which could be especially problematic in vivo, where viral constructs are expressed for weeks, we undertook directed evolution of LATeNT1 in yeast. *S. cerevisiae* yeast are easy to genetically manipulate and libraries of 10^6^-10^8^ cells can be routinely sorted by FACS. To convert LATeNT-catalyzed VAMP2 cleavage into a FACS-detectable signal, we tethered VAMP2 to the yeast plasma membrane and fused its N-terminal end to the chimeric transcription factor LexA-VP16. VAMP2 cleavage frees LexA-VP16 for translocation to the nucleus, where it can drive expression of a YFP reporter (**Fig. 1F**).

We created a library of LATeNT1 variants by error prone PCR and fused to mCherry to enable quantification of expression levels. We first performed positive selection by exposing yeast cultures to room light during overnight induction, then waiting 8 hours for YFP reporter expression. We used FACS to isolate cells with high YFP/mCherry ratio. After amplification of sorted cells, we performed a negative selection without light treatment, and used FACS to retain only cells with low YFP/mCherry ratio. In total, 5 rounds of selection were performed: 3 positive selections and 2 negative selections (**Fig. S1C**). Population analysis by analytical flow cytometry showed progressive improvements in both signal and background over the course of the evolution (**Fig. 1G** and **Fig. S1D**).

After the 5^th^ round of selection, we selected 20 enriched clones for further analysis (**Fig. S2A, B**). Clone 16 (C16), which had one mutation (K34R) in the TeNT protease and one mutation (D397N) in the LOV domain (**Fig. 1B**), showed significantly lower background activity than LATeNT1 while maintaining comparable +light activity in yeast (**Fig. S2C**). This trend was recapitulated in HEK293T cells expressing VAMP2 reporter; the light/dark signal ratio was 97.3, compared to 6.7 for LATeNT1, primarily due to a dramatic reduction in dark state leak (**Fig. 1H**). C16 was named LATeNT, and used for subsequent experiments in this study.

### Characterization of LATeNT in HEK293T cells and neuron culture

To compare the catalytic efficiency of LATeNT with that of wild-type TeNT protease, we prepared TeNT- or LATeNT-containing lysates from HEK293T cells and incubated them with VAMP2-containing lysates for various lengths of time. Quantification of VAMP2 cleavage by Western blot shows that LATeNT has ∼36% the activity of wild-type TeNT (**Fig. 2A** and **Fig. S3A**). To validate LATeNT’s mechanism, we made point mutations in its protease active site or in its LOV domain. E233A in the protease domain and C415A in LOV eliminated VAMP2 cleavage activity, while I504E in the LOV domain made LATeNT constitutively active (**Fig. S3B**).

**Figure 2.**
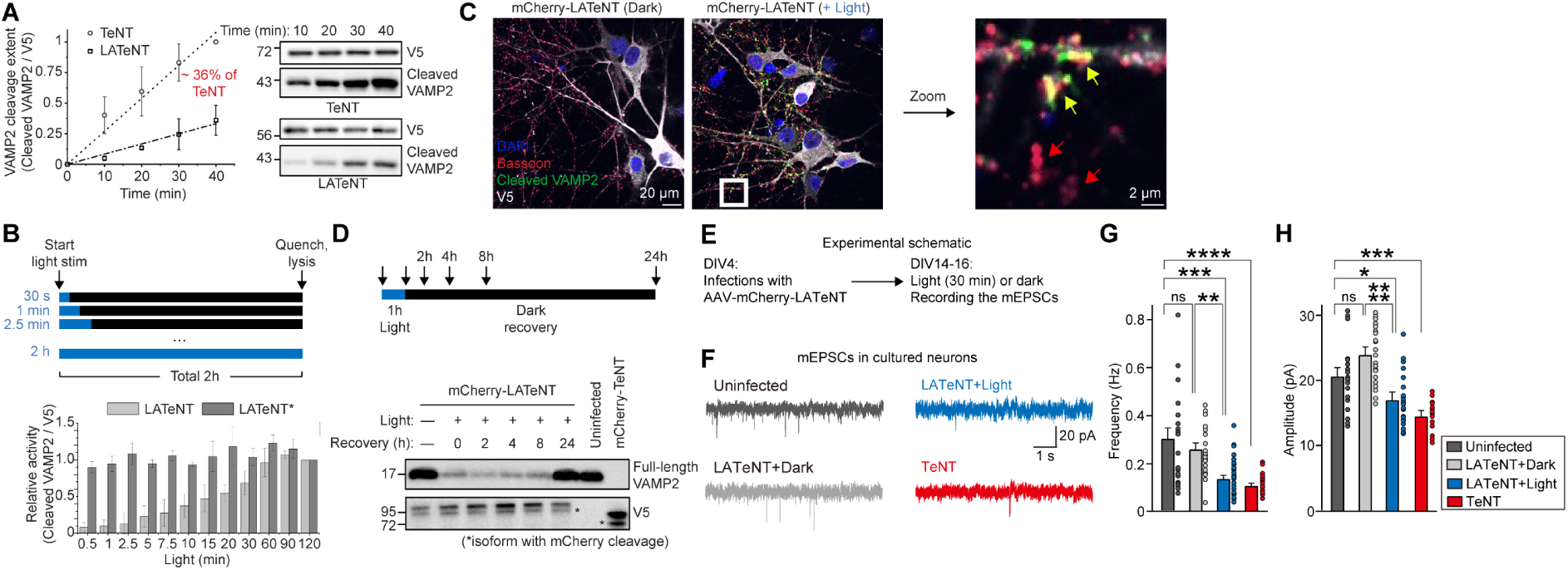
Characterization of LATeNT in HEK293T cells and neuron culture. **(A)** Comparison of TeNT and LATeNT cleavage rates when added to lysate from HEK293T cells expressing recombinant VAMP2. 470 nm light was used. 3 biological replicates; error bars, s.d. **(B)** Kinetics of LATeNT turn-off after light removal. HEK293T cells expressing LATeNT or LATeNT* (V381L) and VAMP2 were stimulated with 470 nm light for 30 sec to 2 h, then maintained in the dark for the remaining time, for a total of 2 hours. 3 biological replicates; error bars, s.d. Representative Western blot data in **Fig. S3E**. **(C)** mCherry-LATeNT cleaves endogenous VAMP2 in cultured neurons. After 30 min stimulation with 470 nm light, rat neurons were fixed and stained with anti-cleaved VAMP2 antibody and anti-Bassoon antibody (pre-synaptic marker). Yellow arrows in zoom show colocalization between Bassoon and cleaved VAMP2 along LATeNT-expressing (white V5-positive) processes. Red arrows point to Bassoon puncta in LATeNT-negative cells lacking cleaved VAMP2 staining. Additional field of views in **Fig. S3F**. **(D)** Endogenous VAMP2 is replenished in neurons after LATeNT-catalyzed cleavage. After LATeNT stimulation with 470 nm light for 1 hour, neurons were allowed to recover for 0-24 hours, then lysates were analyzed by Western blotting. This experiment was performed 3 times with similar results. Additional blots in **Fig. S3G**. **(E)** Scheme for miniature excitatory postsynaptic current (mEPSC) recordings in cultured hippocampal rat neurons. **(F)** Representative traces of mEPSCs recorded from uninfected neurons, LATeNT-expressing neurons kept in the dark (light gray) or exposed to 0.55 mW/cm^2^ LED light for 30 min (blue). **(G, H)** Quantification of frequencies (G) and amplitudes (H) of mEPSCs recorded in (F). Data are presented as mean ± SEM (uninfected, n = 19; LATeNT+Dark, n = 21; LATeNT+Light, n = 26; TeNT, n = 15; *p < 0.05, **p < 0.01, ***p < 0.001, ****p < 0.0001; ANOVA with Tukey’s *post hoc* test).

Next we characterized LATeNT’s light requirements. Continuous 470 nm blue light at 1 mW/cm^2^ power gives maximal LATeNT activation (**Fig. S3C**). However, 2 second pulses with the same light power (16% duty cycle; 2 sec on/10 sec off per cycle) gave a comparable degree of activation (**Fig. S3D**). These mild conditions use much less light power than the >10 mW/cm^2^ light intensity typically used to activate halorhodopsins^28, 29^.

To investigate LATeNT’s reversibility, we performed an experiment in which LATeNT was activated with light for 30 sec to 2 h, then incubated in the dark for the remainder of the 2 h experiment (**Fig. 2B**). We observed a linear increase in VAMP2 cleavage with light exposure time, independent of the dark incubation time (**Fig. 2B** and **Fig. S3E**), suggesting that LATeNT activity shuts off rapidly, within a few minutes of light removal. This is similar to observations with other LOV-based tools^23, 30^. We also made a point mutant of LATeNT, named LATeNT*, carrying a V381L mutation in LOV that is known to slow LOV domain closing (τ = 55 sec for wild-type LOV versus 72 min for V381L-LOV^31^). VAMP2 cleavage by LATeNT* reached saturation regardless of illumination time or dark reset time (**Fig. 2B** and **Fig. S3E**), indicating that LATeNT* remains active for over 60 min even after brief illumination for <1 minute.

In neurons, endogenous VAMP2 is primarily localized in presynaptic vesicles at axonal termini^32, 33^. To determine whether LATeNT can traffic through axonal processes and cleave endogenous VAMP2, we expressed LATeNT in dissociated cortical rat neuron cultures using adeno-associated virus (AAV) infection. Confocal imaging of stained neurons shows the presence of LATeNT along processes and at synaptic termini positive for the marker Basson (**Fig. 2C**). Upon light stimulation, processes expressing LATeNT displayed cleaved VAMP2 puncta colocalized with Bassoon (**Fig. 2C**, yellow arrows), whereas processes lacking LATeNT showed no cleaved VAMP2 signal (**Fig. 2C**, red arrows). Interestingly, in control experiments with wild-type TeNT, cleaved VAMP2 signal was detected primarily in the cell body (**Fig. S3F**), probably because immediate cleavage of VAMP2 after its translation removes its synaptic vesicle targeting domain^34^. Western blot analysis of cell lysates from cultured neurons showed that 1 hour of light exposure depleted endogenous VAMP2 levels by 79.3 ± 2.5% (**Fig. 2D** and **Fig. S3G**), which was fully restored after ∼24 hours of recovery in the dark (**Fig. 2D**).

Finally, to assess the functional effects of LATeNT activity in neurons, we transduced dissociated hippocampal neuron cultures and measured miniature excitatory postsynaptic currents (mEPSCs) (**Fig. 2E**). Exposure of LATeNT-expressing neurons to light led to a significant reduction in both the frequency and amplitude of mEPSCs, comparable to the effects observed with wild-type TeNT (**Fig. 2F-H**).

### LATeNT inhibits synaptic transmission in the mouse brain

To characterize LATeNT in the mouse brain, we injected AAVs encoding mCherry-LATeNT into the hippocampal CA3 region of adult mice and prepared acute brain slices two weeks post-injection (**Fig. 3A** and **Fig. S4A**). We measured synchronous evoked EPSCs (eEPSCs) in hippocampal CA1 pyramidal neurons, evoked by simulating axon fibers of the Schaffer-collateral (SC) pathway, both in the absence and presence of light.

**Figure 3.**
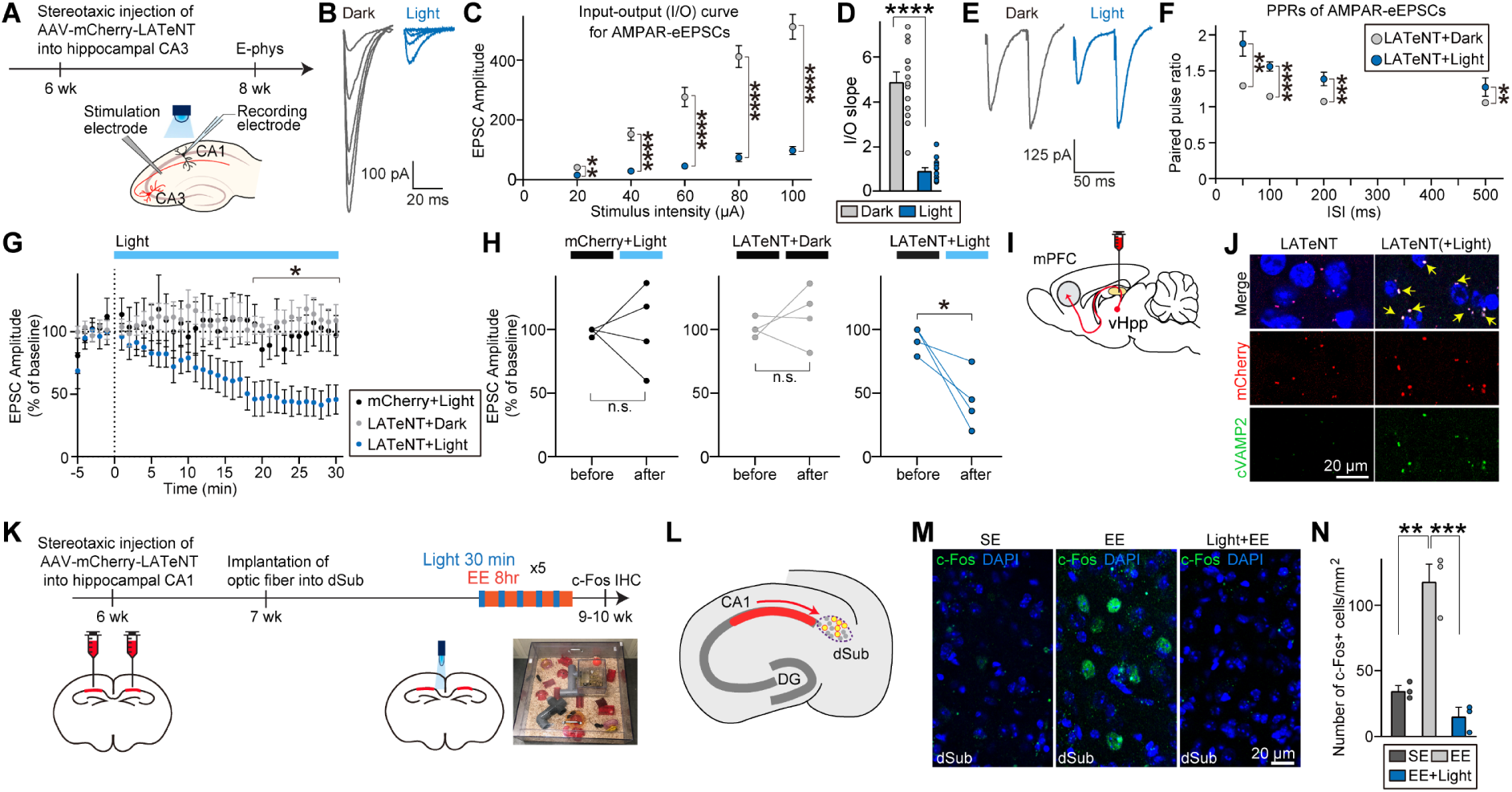
LATeNT inhibits synaptic transmission in the mouse brain. **(A)** Experimental scheme. Synaptic connectivity was tested by stimulating presynaptic fibers from hippocampal CA3 and monitoring postsynaptic responses in CA1 pyramidal neurons. Light treatment was ambient room light (0.15∼0.2 mW/cm^2^) for 1∼2 h. **(B-D)** Excitatory synaptic strength, measured using input-output (I-O) curves, from CA3-CA1 synapses. Representative AMPAR-EPSC traces (B), summary plots of the EPSC amplitudes as a function of CA3 stimulation current (C), and a summary graph of fitted linear I-O slopes (D). Data are presented as mean ± SEM (‘n’ denotes the number of recorded neurons/mice; dark, n = 14/4; light, n = 14/4; Mann-Whitney *U* test). **(E, F)** EPSC-PPRs (paired pulse ratios) at CA3-CA1 synapses as a function of interstimulus interval. Representative trace of EPSC-PPRs in (E) at an interstimulus interval of 50 ms. Data in (F) are presented as mean ± SEM (‘n’ denotes the number of recorded neurons/mice; dark, n = 13/4; light, n = 12/4; Mann-Whitney *U* test). **(G)** EPSC recording during continuous light illumination of mCherry-LATeNT-expressing brain slices. 473-nm blue light was provided at 0.5 mW/cm². Data are presented as mean ± SEM (‘n’ denotes the number of recorded neurons/mice; mCherry+Light, n = 4/3; LATeNT+Dark, n = 4/3, LATeNT+Light, n = 4/3; two-way repeated measures ANOVA with Tukey’s *post hoc* test). **(H)** Paired plots of EPSC amplitudes averaged over the first 5 min (before) and last 5 min (after). Each pair of line-connected dots represents an individual neuron (*p < 0.05; paired t-test). **(I)** Experimental scheme for testing LATeNT in long-range projections of adult mice. AAVs were injected into the vHPP and an optical fiber was implanted into the mPFC. Light (473 nm) was delivered to the mPFC through the optical fiber for 30 min (1.0 mW/cm2, 2s on, 10s off). **(J)** Representative images of mPFC regions stained with anti-cleaved VAMP2 (green) antibody, from experiment in (I). Yellow arrows indicate colocalization of mCherry with cleaved VAMP2 puncta. **(K)** Experimental scheme for testing LATeNT effect on c-Fos expression after exposure to enriched environments (EE). AAVs were injected into hippocampal CA1, and an optical fiber was implanted into the dorsal subiculum (dSub). 3 weeks after injection, 473 nm light was delivered to the dSub for 30 min (1.0 mW/cm^2^, 2s on, 10s off), after which mice were provided EE conditions for 8 h per day for 5 consecutive days. Sections were stained with anti-c-Fos antibody. **(L)** Hippocampal CA1 neurons expressing mCherry-LATeNT (red) projecting into dSub. **(M)** Representative images of c-Fos immunostaining in the dSub region. SE, standard environment. Scale bar = 20 μm. **(N)** Quantitative analysis of the number of c-Fos–expressing cells per unit area in mice under SE (dark gray), EE (light gray), or EE+Light (blue) conditions. Data are presented as mean ± SEM (n = 3 mice each after averaging data from 4 sections/mouse; **p < 0.01, ***p < 0.001; ANOVA with Tukey’s *post hoc* test).

Exposure of LATeNT-expressing slices to light prior to recordings resulted in a significant reduction in the amplitudes of AMPAR-mediated eEPSCs compared to slices kept in the dark, as assessed by input-output (I-O) curves (**Fig. 3B-D**). Similarly, NMDAR-mediated responses also showed a decrease in synaptic strength, as indicated by a comparable NMDAR/AMPAR ratio in both light- and dark-conditioned neurons (**Fig. S4B-C**), suggesting that LATeNT inhibited both AMPAR and NMDAR responses. Furthermore, the reduction in excitatory synaptic strength in the SC pathway was accompanied by a significant decrease in neurotransmitter release probability (P_r_), as evidenced by an increased paired-pulse ratio (PPR) following light stimulation (**Fig. 3E-F**).

To investigate the real-time functional kinetics of LATeNT, we measured AMPAR-mediated eEPSCs every 1 min in hippocampal CA1 pyramidal neurons obtained from mice injected with AAVs expressing mCherry-LATeNT or mCherry in the hippocampal CA3 region. During repeated measurements, baseline amplitudes were first established in the dark, followed by continuous light illumination (0.5 mW/cm^2^) of brain slices. In LATeNT-expressing neurons, eEPSC amplitudes gradually decreased upon light illumination, reaching statistical significance after 20 minutes, whereas no changes were observed in groups without LATeNT or kept in the dark (**Fig. 3G**). This result was further validated by paired comparison of EPSC amplitudes from individual cells (**Fig. 3H**).

To test whether LATeNT can control neurotransmission in long-range projections, we injected AAVs expressing mCherry-LATeNT into the ventral hippocampus (vHPP) of adult mice; this strategy was chosen because vHPP neurons project to the prelimbic (PL) regions of the medial prefrontal cortex (mPFC)^35^ (**Fig. 3I** and **Fig. S4D**). Three weeks later, mice were exposed to blue light for 30 min via a fiber implanted in the mPFC. Quantitative immunofluorescence analysis revealed a light-dependent increase in the colocalization of cleaved VAMP2 puncta with mCherry puncta (**Fig. 3J** and **Fig. S4E**), indicating that LATeNT trafficks to the synaptic termini of mPFC-projecting vHPP neurons.

Finally, to test LATeNT *in vivo* in the context of behavior, we used an enriched environment (EE) protocol, which has been shown to enhance neuronal plasticity and activity in response to experience^36^. LATeNT AAVs were injected into the hippocampal CA1 of both hemispheres of adult mice, which were then housed in either a standard environment (SE) or EE (**Fig. 3K-L, Fig. S4F**). In the EE condition, mice were either kept in the dark or treated with light (delivered to the dorsal subiculum (dSub) region through an implanted fiber), then placed in EE for 8 hours, repeated for a total of five sessions (**Fig. 3K**). As expected, mice housed in EE showed a robust increase in c-Fos expression in dSub compared to mice housed in SE (**Fig. 3M-N**). However, this increase was completely absent in mice that received light prior to EE (**Fig. 3M-N**), suggesting that LATeNT activation inhibited synaptic transmissions from CA1 to dSub, which is necessary for elevated c-Fos expression. The light-dependent activation of LATeNT was further confirmed by staining for cleaved VAMP2 (**Fig. S4G-H**). As a control, light stimulation did not affect the EE-dependent increase in c-Fos expression in the auditory cortex (AC) (**Fig. S4I-J**), which does not receive direct synaptic inputs from hippocampal CA1. These results suggest that LATeNT is an effective tool for spatiotemporal manipulation of neuronal activity *in vivo*.

### LATeNT reveals a causal role for CA1 SST interneurons in regulating anxiety-like behaviors in mice

Having established the utility of LATeNT *in vivo*, we sought to apply it to understand the functional relationship between a specific hippocampal cell population and anxiety-like behaviors. Previously, our lab discovered that conditional knockout of NPAS4, an activity-dependent transcription factor controlling GABAergic synapse development, in hippocampal CA1 SST^+^ (somatostatin-positive) interneurons increases anxiety-like behaviors^37^. However, as NPAS4 affects multiple pathways, including development^38^, it is unclear which specific process underlies the observed changes in anxiety behavior. By using LATeNT to precisely inhibit synaptic transmission in CA1 SST^+^ interneurons, we wished to test the hypothesis that this microcircuit regulates anxiety-like behavior in mammals.

We expressed LATeNT selectively in SST^+^ GABAergic interneurons in the mouse hippocampal CA1 by injecting Flp-dependent AAVs into the hippocampal CA1 region of adult *Sst*-IRES-FlpO mice. The viruses expressed either mCherry-LATeNT or mCherry alone as a control (**Fig. 4A** and **4B**). Immunohistochemical analysis with anti-SST antibodies confirmed that mCherry fluorescence was specifically localized to SST^+^ interneurons, validating the targeted expression of the construct (**Fig. 4C**).

**Figure 4.**
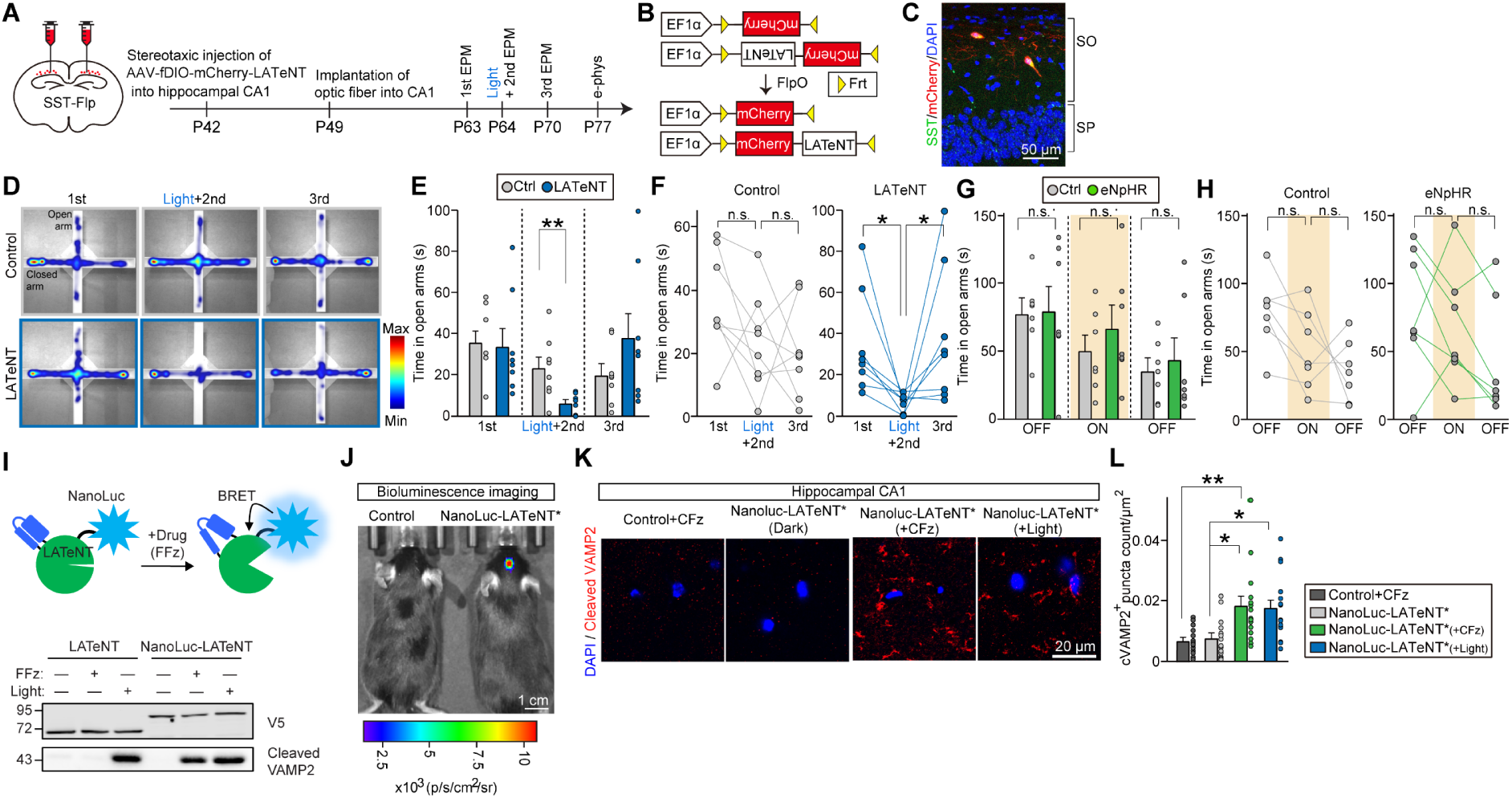
LATeNT identifies causal role for CA1 SST^+^ interneurons in anxiety-like behaviors. **(A)** Experimental schematic. AAVs expressing fDIO-mCherry-LATeNT or fDIO- mCherry were injected into the hippocampal CA1 of Sst-IRES-FlpO mice, and optical fibers were implanted into the hippocampal CA1. Three weeks after injection, the first elevated plus maze (EPM) test was performed. The next day, the second EPM test was performed 6 hours after 473 nm light was delivered to the CA1 through an optical fiber for 30 min (2 sec on/10 sec off). The third EPM test was performed one week later. **(B)** FlpO-mediated recombination of AAV-delivered genes. **(C)** Representative image showing mCherry-LATeNT expression in hippocampal CA1 SST interneurons. SO, stratum oriens, SP, stratum pyramidale. Scale bar = 50 μm. **(D)** Representative heatmaps of time spent in open versus closed arms of the EPM. **(E)** Bar graphs presenting the time spent in open arms across trials. Data are presented as mean ± SEM (n = 8 mice; *p < 0.05; Mann-Whitney *U* test). **(F)** Before and after graphs presenting the time spent in open arms across trials. Each pair of line-connected dots represents an individual mouse. 6-8 mice analyzed per condition (*p < 0.05; Friedman test). **(G)** Same experiment as (A) except the halorhodopsin eNpHR3.0^40^ was used to inhibit neuronal activity instead of LATeNT (detailed schematic in **Fig. S5I**). Bar graphs show the time spent in open arms across trials for eNpHR-expressing mice versus non-expressing controls. Data are presented as mean ± SEM (n = 7 mice; *p < 0.05; Mann-Whitney *U* test). **(H)** Before and after graphs presenting the time spent in open arms across trials. Each pair of line- connected dots represents an individual mouse. 7 mice were analyzed per condition (Friedman test). **(I)** LATeNT fusion to the luciferase NanoLuc allows LATeNT to be activated by a small-molecule (fluorofurimazine, FFz) instead of light. HEK293T cells expressing NanoLuc-LATeNT were treated with blue light for 30 min or FFz for 60 min. This experiment was performed 3 times with similar results. **(J-L)** NanoLuc-LATeNT* can be activated by cephalofurimazine (CFz) *in vivo* (detailed schematic in **Fig. S7**). (J) Representative bioluminescence images of mice expressing NanoLuc-LATeNT* in the hippocampus and injected with CFz. Control mice do not express LATeNT*. (K) Representative images of hippocampal CA1 regions stained with anti-cleaved VAMP2 (green) antibody. (L) Summary graphs quantifying cVAMP2^+^ puncta density in (K). Data are presented as mean ± SEM (n = 15 brain sections from 3 mice per group; *p < 0.05, **p < 0.01; ANOVA with Tukey’s *post hoc* comparisons test).

Three weeks post-injection, we monitored anxiety-like behaviors in these mice by performing an elevated plus maze (EPM) test. In the 1^st^ EPM test, performed in the absence of light stimulation, both mCherry- and mCherry-LATeNT-expressing mice exhibited comparable anxiety-like behaviors, spending a similar amount of time in open arms (1^st^ EPM test; **Fig. 4D- E**). In marked contrast, following blue light illumination via CA1-implanted fibers, mCherry- LATeNT-expressing mice showed increased anxiety-like behaviors compared with light-exposed control mice, exhibiting a significant decrease in time spent in the open arms (2^nd^ EPM test; **Fig. 4D-F**). To determine whether the increased anxiety-like behaviors induced by inhibiting the activities of CA1 SST^+^ interneurons could be reversed, we performed a 3^rd^ EPM test on the same mice 1 week later – sufficient time to allow full recovery of VAMP2 (see **Fig. 2D**). We observed that the light-induced increased anxiety-like behavior (decrease in time spent in open arms) was fully reversed at this time point, with mCherry-LATeNT-expressing mice behaving comparably to control mice (3^rd^ EPM test; **Fig. 4D-F**).

Hippocampal CA1 SST^+^ interneurons innervate the distal dendritic compartment of pyramidal neurons in the CA1 hippocampus^39^ (**Fig. S5A**). To validate the LATeNT-mediated inhibition of SST^+^ interneuron activity, we stimulated axons in *stratum lacunosum-moleculare* (SLM; **Fig. S5A**) and *stratum pyramidale* (SP; **Fig. S5E**) layers under ambient room light and measured dendritic and somatic eIPSCs, respectively, by performing whole-cell patch clamp recordings. The amplitude of dendritic, but not somatic, eIPSCs was decreased in hippocampal CA1 pyramidal neurons of LATeNT-expressing mice compared with mCherry-expressing mice (**Fig. S5B-C** versus **Fig. S5F-G**). Moreover, dendritic, but not somatic, eIPSC-PPRs were significantly increased in LATeNT-expressing mice, indicating decreased neurotransmitter release probability in CA1 SST^+^ interneurons **(Fig. S5D** versus **Fig. S5H)**. These results are consistent with LATeNT acting in a light-dependent manner to suppress neurotransmitter release from SST^+^ interneurons onto CA1 pyramidal neurons.

Several light-driven ion channels and pumps, such as enhanced halorhodopsin (eNpHR3.0^40^), have been widely used to dissect neural circuitry due to their ability to control neuronal spiking on a millisecond timescale. However, such tools are limited when silencing is required for extended periods of time, as sustained light illumination results in tissue heating and phototoxicity. To test whether acute, halorhodopsin-mediated silencing of CA1 SST^+^ interneurons could also elicit similar changes in anxiety-like behaviors, we repeated the same experiment using eNpHR. We stereotactically injected AAV-DIO-eNpHR-EYFP or AAV-DIO-EYFP (control) into the hippocampal CA1 of adult *Sst*-IRES-Cre mice to achieve specific expression at SST^+^ interneurons (**Fig. S5I-K**). At 3 weeks post-injection, we performed EPM tests on a single 9-min session comprising the following three 3-min epochs: a light-off baseline epoch, a light-on illumination epoch with constant illumination of 594 nm light (5 mW), and a second OFF epoch.

Unlike the LATeNT-mediated inhibition, the eNpHR-mediated inhibition of CA1 SST^+^ interneurons did not induce any alteration of the time spent in open arms during all three sessions (**Fig. 4G-H, Fig. S5L)**. We validated eNpHR-mediated inhibition of CA1 SST^+^ interneurons by measuring firing rates, and found that neurons expressing eNpHR exhibited decreased firing rates under yellow-light illumination of brain slices (**Fig. S5M-N**). These results suggest that inhibition of CA1 SST^+^ interneuron-mediated anxiety-like behaviors requires a longer silencing period, which can be achieved *in vivo* by LATeNT but not by eNpHR.

These experiments establish a causal link between GABAergic synaptic transmission in CA1 SST^+^ interneurons and anxiety-like behaviors, made possible by LATeNT’s unique ability to sustainably and reversibly inhibit the activity of defined neural populations *in vivo*.

### NanoLuc-LATeNT for drug-activated VAMP2 cleavage

The use of light to control LATeNT activation has the benefit of spatial and temporal precision. However, light delivery to the mouse brain requires implantation of an optical fiber and is restricted to small regions. A drug-controlled version of TeNT could be easier to use, across multiple brain regions at once if desired. To enable this, we made use a previous observation that LOV-containing optogenetic tools can be activated by bioluminescence resonance energy transfer (BRET) from a blue light-emitting luciferase instead of external light^41, 42^. To explore this for LATeNT, we fused it directly to the luciferase NanoLuc, which emits 460 nm blue light (**Fig 4I**, top). We expressed the construct in the cytosol of HEK293T cells along with VAMP2 reporter. **Fig. 4I** shows that cells treated with NanoLuc’s small-molecule substrate, fluorofurimazine (FFz), for 60 min in the dark produced a similar degree of VAMP2 cleavage as cells exposed to blue light for 30 min.

We then tested the possibility of uncaging LATeNT activity with drug *in vivo*. We stereotactically injected AAVs expressing NanoLuc-LATeNT* into the hippocampal CA3 region of adult mice and performed intraperitoneal injections of cephalofurimazine (CFz)^43^, a NanoLuc substrate optimized for *in vivo* use (**Fig. S6A**). Functional expression of NanoLuc-LATeNT* was confirmed by in vivo bioluminescence imaging (**Fig. 4J**). When tissue sections were analyzed by immunofluorescence staining, we observed cleavage of endogenous VAMP2 in mice expressing NanoLuc-LATeNT* and treated with CFz, but not in controls lacking LATeNT* or CFz (**Fig. 4K-L**). Interestingly, the increase in cleaved VAMP2 was comparable to that observed in mice subjected to light via an implanted optical fiber. These results suggest that NanoLuc-LATeNT* is a promising alternative for drug- rather than light-controlled turn-on of VAMP2 cleavage and synaptic inhibition *in vivo*.

## Control of endogenous insulin secretion using LATeNT

VAMP2 is expressed predominantly in the brain, but a few other professional secretory cells use VAMP2 as well^18^, such as pancreatic beta cells that secrete insulin. The ability to control endogenous insulin release could have benefits for diseases in which insulin is dysregulated, such as insulinomas and hyperinsulinemia^44^. Because insulin exocytosis from pancreatic beta cells requires VAMP2, we investigated whether LATeNT could be used to regulate this process.

We expressed LATeNT in MIN6 cells, a well-established model of glucose-stimulated insulin secretion (GSIS) from beta cells^45^. Cells were then starved in low glucose for 1 hour, then stimulated with high glucose for 1 hr to induce GSIS (**Fig. 5A**). In cells lacking LATeNT, high glucose increased GSIS by 2.0- to 2.3-fold, consistent with previous reports^45^ (**Fig. 5A**). LATeNT-expressing cells, when kept in the dark, showed a similar GSIS response (2.4-fold) (**Fig. 5A**). However, GSIS was impaired when LATeNT-expressing cells were stimulated with light for 30 minutes prior to glucose starvation (**Fig. 5A**). Western blotting and confocal imaging confirmed LATeNT- and light-dependent cleavage of endogenous VAMP2. (**Fig. S6B-C**). This result demonstrates that LATeNT can be deployed to regulate other endogenous processes of biological and therapeutic importance, in non-neuronal cells.

**Figure 5.**
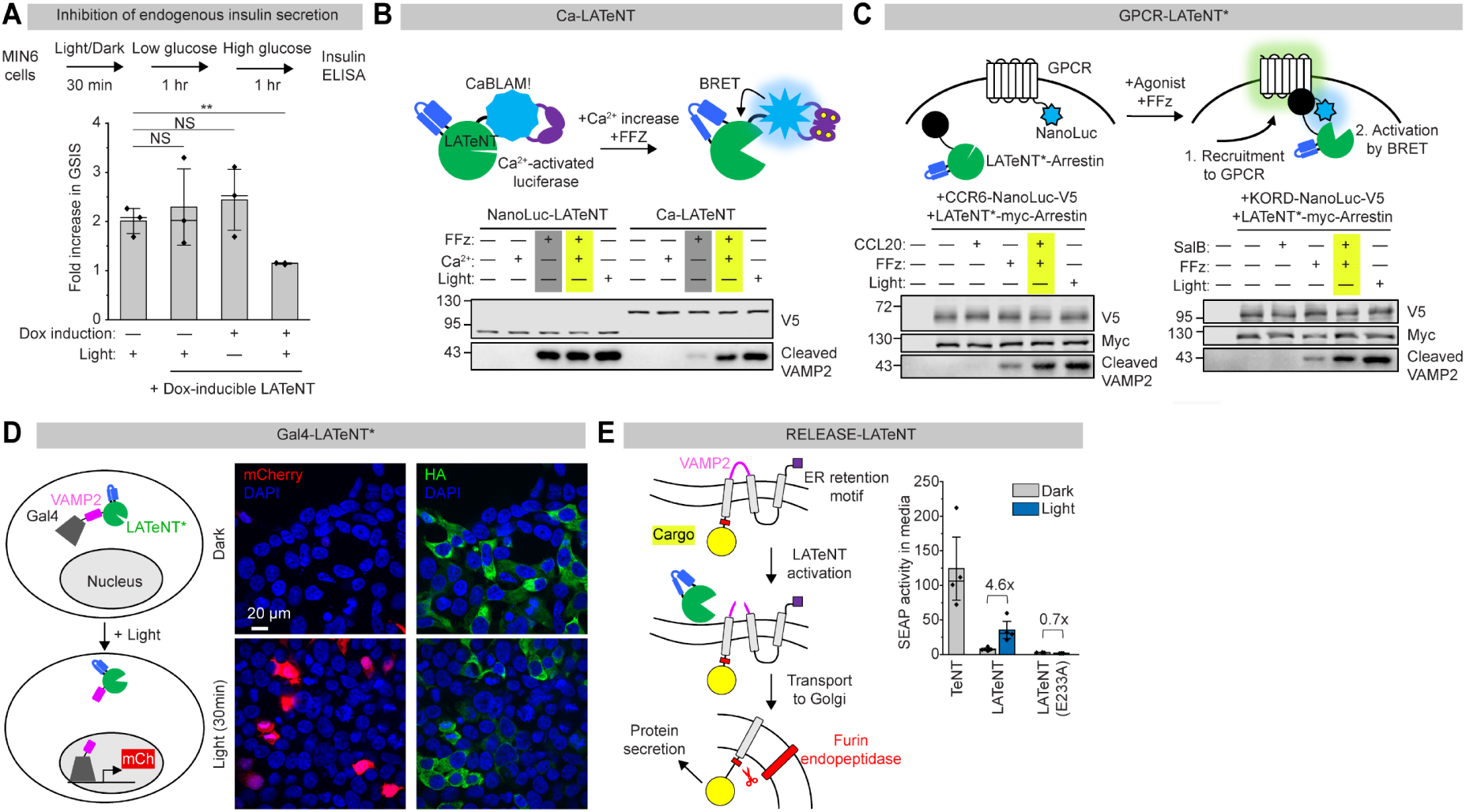
LATeNT for regulation of insulin secretion and construction of synthetic circuits. **(A)** LATeNT control of endogenous insulin secretion. Top: experimental schematic; switching from low to high glucose stimulates insulin secretion in MIN6 pancreatic beta cells. Bottom: Fold increase in glucose-stimulated insulin secretion (GSIS) before/after doxycycline induction of LATeNT expression and blue light exposure for 30 minutes (prior to low glucose treatment). 3 biological replicates; error bars, s.d.. **p < 0.01, Student’s t-test. **(B)** Calcium activated LATeNT through fusion of LATeNT to the Ca^2+^-activated luciferase CaBLAM!^47^. Fluorofurimazine (FFz) is used by CaBLAM! to generate bioluminescence. This experiment was performed 3 times with similar results. **(C)** GPCR-LATeNT* converts GPCR activity into LATeNT* turn-on. HEK293T cells expressing GPCR-NanoLuc and LATeNT*-arrestin were stimulated with agonist (0.2 μg/mL CCL20 or 1 μM Salvinorin B) and FFz for 60 minutes, or light for 30 minutes. CCR6, C-C chemokine receptor 6. KORD, κ-opioid receptor DREADD. This experiment was performed 3 times with similar results. **(D)** Gal4-LATeNT* is a single- component, light-dependent transcription factor. Anti-HA stain shows cytosolic localization of the construct. mCherry expression requires light. This experiment was performed 3 times with similar results. **(E)** RELEASE-LATeNT enables light-triggered secretion of protein cargoes. LATeNT activity separates the cargo from the ER retention motif, allowing cargo trafficking to the cell surface. HEK293T cells expressing RELEASE-LATeNT constructs were stimulated with 470 nm light for 30 min and released SEAP (alkaline phosphatase) reporter activity in the media was measured 4 h later. 4 biological replicates; error bars, s.d.

### Synthetic biology applications of LATeNT

Proteases are increasingly used as building blocks in the construction of synthetic circuits, allowing faster responses and computations than gene transcription-based circuits^46^. For such applications, sequence-specific orthogonal proteases are required. We wondered if LATeNT could be used for the construction of synthetic circuits in non-neuronal cells where VAMP2 expression is absent.

First, we engineered LATeNT to respond to different inputs. Calcium is a ubiquitous second messenger used in immune cell activation, cell proliferation, apoptosis, muscle contraction, neurotransmitter release, and numerous other signaling processes. To engineer LATeNT to respond to elevated cytosolic calcium, we fused it to CaBLAM!^47^, a recently-reported Ca^2+^- dependent luciferase. **Fig. 5B** shows that “Ca-LATeNT” expressed in HEK293T cells gave 6.7- fold greater cleavage of VAMP2 reporter in the presence of Ca^2+^ and fluorofurimazine (FFz, used by luciferase to generate bioluminescence) than when treated with FFz alone. 60-minute exposure to Ca^2+^/FFz produced a similar extent of VAMP2 cleavage as 30 min treatment with light, indicating that CaBLAM! in the high Ca^2+^ state provides efficient BRET-activation of LATeNT.

Next, we engineered LATeNT to be turned on by the activation of a G-protein coupled receptor (GPCR). We used the chemokine GPCR CCR6, which is activated by its peptide ligand CCL20^48^. We expressed CCR6-NanoLuc with LATeNT*-arrestin in HEK293T cells, and added CCL20 to stimulate LATeNT*-arrestin recruitment to the receptor. The resulting proximity between LATeNT* and CCR6-fused Nanoluc enables BRET activation of the former, if FFz is also present (**Fig. 5C**, top). We observed 2.3-fold higher VAMP2 cleavage in the presence of CCL20 than in its absence, and no VAMP2 cleavage was detected in the absence of FFz (**Fig. 5C**, bottom). We performed a similar experiment with a different GPCR, the kappa opioid receptor-based DREADD (KORD)^49^, which is activated by the drug salvinorin B (SalB). We again observed 2.1-fold higher VAMP2 cleavage in the presence of SalB than in its absence (**Fig. 5C**, bottom). These examples show that LATeNT can be gated by diverse inputs, including Ca^2+^ and receptor activity.

We then explored LATeNT’s ability to trigger different outputs besides synaptic inhibition. To this end, we created LATeNT variants that drive either transgene expression or protein secretion. For LATeNT-induced gene expression, we created a fusion between Gal4, VAMP2, and LATeNT*, and expressed the construct in HEK293T cells along with a UAS-mCherry reporter (**Fig. 5D**). This construct localizes to the cytosol in the absence of light due to its large size (112 kDa). Upon light stimulation, intramolecular cleavage of the VAMP2 domain releases the smaller Gal4 fragment (38 kD) to translocate to the nucleus and drive gene expression (**Fig. 5D**). Confocal imaging shows mCherry reporter expression only in cells exposed to light, and not in dark-maintained cells (**Fig. 5D**).

For a more immediate, non-transcriptional response to LATeNT activation, we designed a circuit using the RELEASE^50^ tool which uses protease activity to control protein secretion from the endoplasmic reticulum (ER). Our design in **Fig. 5E** uses the cytosolic domain of VAMP2 as the protease cleavage sequence and secreted alkaline phosphatase (SEAP) as the cargo. Proteolytic cleavage of our construct separates the cargo from a C-terminal ER retention motif, enabling the cargo to traffic through the trans-Golgi network into the extracellular medium. 4 hours after LATeNT and RELEASE-expressing cells were stimulated light, we detected 4.6- fold higher SEAP activity in the media than in control cultures not exposed to light (**Fig. 5E**). Mutation of LATeNT’s active site (E233A) to abolish protease activity suppressed all SEAP secretion, even after light exposure (**Fig. 5E**).

Together, our results demonstrate that LATeNT can be a versatile tool to convert various biologically-relevant inputs into gene expression or protein secretion in living mammalian cells.

## Discussion

LATeNT is an optogenetic tool that silences neurons through light-dependent proteolytic cleavage of the synaptic vesicle fusion protein VAMP2. By alternating positive and negative selections during directed evolution, we were able to largely retain the ‘on’ state activity of wild-type TeNT while suppressing dark state background to nearly undetectable levels. LATeNT efficiently silenced synaptic transmission upon blue light exposure in vivo, leading to the discovery of a causal link between synaptic transmission in a specific interneuron population and anxiety-like behavior in mice. LATeNT-mediated VAMP2 cleavage enabled manipulation of other endogenous processes, such as insulin secretion in pancreatic beta cells. Finally, LATeNT’s versatility in synthetic biology applications was shown by creating variants that respond to different inputs (small-molecule drug, Ca^2+^, or receptor activity) to produce diverse outputs (gene expression and protein secretion).

As a neuronal silencing tool, LATeNT fills an unmet need by providing rapidly-inducible, sustained, and reversible silencing of synaptic transmission. Such manipulation is required in systems neuroscience to establish causal relationships between defined cell populations and functional readouts or behavior. Existing tools for long-term silencing (hours to days) include DREADDs, constitutively-active TeNT protease, and overexpression of inwardly rectifying potassium (Kir) channels^51, 52^. However, DREADDs are weak and affect multiple pathways downstream of G protein activation^51, 53^ including β-arrestin, ERK, adenyl cyclase, cAMP, and Ca^2+^/Na^+^ currents in addition to neurotransmitter release. Long-term expression of wild-type TeNT and Kir can produce developmental defects^54^.

The predominant tool for short-term neuronal silencing is halorhodopsin. However, opsins are limited when silencing is required for extended periods of time, as sustained light illumination results in tissue heating and phototoxicity. Furthermore, we showed that the halorhodopsin eNpHR is insufficiently potent to induce a behavioral phenotype in the context of SST^+^ interneuron modulation of anxiety, whereas LATeNT’s sustained action produced a robust and reproducible effect. Of note, a separate light-regulated neurotoxin, PA-BoNT, has been reported for in vivo modulation of neuronal activity^55^, but in a side by side comparison to LATeNT, PA-BoNT showed much lower +light activity and higher dark-state background than LATeNT (**Fig. S7A**).

In addition to LATeNT, our study presents LATeNT*, a V381L point mutant of LATeNT that reverses much more slowly from the “on state” back to the dark/off state. Even brief illumination for just 30 seconds allowed LATeNT* to continue cleaving VAMP2 for 60 min or longer (**Fig. 2B**). LATeNT* provides a useful alternative in situations requiring maximal effect with minimal light exposure.

LATeNT provides an example, along with several recently-reported tools^22, 23, 56, 57^, of how conformation-switching domains (LOV, CaM-M13 module, etc.) can be used to regulate enzyme activity through engineered allostery. Both LATeNT and our previously-developed LOV-Turbo^23^ exhibit outstanding dynamic range, with negligible activity in the dark state, and activity close to the parental enzyme in the light state (36% in the case of LATeNT, **Fig. 2A**). To explore the generality of LATeNT’s design, we also inserted LOV into the analogous location of botulinum toxin serotype A (BoNT/A) light chain protease, which has moderate sequence similarity (50%) but high structural similarity to TeNT (**Fig. S7B**)^58^. With minimal linker optimization, light-activated BoNT/A (LABoNT/A) gave a 4.4-fold increase in cleavage of its natural substrate SNAP25 upon light exposure (**Fig. S7C**). Since BoNT protease has been re-engineered to cleave non-native substrates^17^, it could be an even more versatile platform than TeNT for the development of regulatable protease tools for precision protein manipulation.

Several lines of future work could improve the utility and scope of LATeNT. First, engineering LATeNT to cleave proteins other than VAMP2 could expand the space of possible applications. Discovering a peptide recognition sequence for LATeNT/TeNT (in place of VAMP2) would also be helpful for creating more compact tools. Second, given LATeNT’s promise in controlling endogenous insulin secretion, it is worth exploring LATeNT for more complex applications – such as control of other hormones including glucagon secretion from pancreatic alpha cells^59^, and sense-and-respond technologies that integrate wearable, real-time glucose sensors^60^. Third, if LATeNT activity can be regulated by cell-cell contact, then it could be a powerful way to manipulate the activity of defined synaptic connections in the brain.

## Acknowledgements

This work was supported by the Chan Zuckerberg Biohub - San Francisco, the Wu Tsai Neurosciences Institute through the Neuro-omics Initiative (to A.Y.T.), and NSF Neuronex (to A. Y. T.). In addition, we are grateful for support from the National Research Foundation of Korea (NRF) funded by the Ministry of Science and ICT (RS-2023-00207834, RS-2022- NR067821, and RS-2023-NR076948 to J.W.U.). Anti-cleaved VAMP2 antibodies were kind gifts from Brigitte Dorner (Robert Koch Institute). Rat cortical tissue was a kind gift from Sungmoo Lee and Michael Lin (Stanford University). MIN6 cell line was a kind gift from Mohammad Azizzanjani and Peter Jackson (Stanford University). Anti-Bassoon antibody was a kind gift from Min Huang and Thomas Südhof (Stanford University). We thank Junhao Xu (Stanford University) for assistance with Ca-LATeNT experiments, and Yaereen Dho (Stanford University) for assistance with preparing CFz solid formulations. A.Y.T. is a Chan Zuckerberg Biohub – San Francisco investigator.

## Author contributions

H.R. and A.Y.T. conceived this project. H.R., J.W.U., and A.Y.T. designed experiments and analyzed the data. H.R. performed all LATeNT engineering, characterization, synthetic biology applications, and endogenous VAMP2 cleavage assay in neurons, with the assistance of C.L.

D.K. carried out stereotactic injections, optic fiber implantations, immunohistochemical analyses, and behavioral analyses in mice. B.K., Y.J., and Y.K. performed electrophysiological analyses. M. J. provided preliminary data on LOV insertion sites. F.X. performed LABoNT/A experiments. H.R., J.W.U., and A.Y.T. wrote the manuscript with input from all authors.

## Methods

### Cloning

For LATeNT constructs, gene fragments were ordered from Twist Biosciences and were subcloned into a pcDNA3 backbone (https://www.addgene.org/vector-database/2093/) using HindIII and XhoI restriction sites. All constructs were generated using standard cloning techniques. PCR fragments were amplified using Q5 polymerase (NEB Cat# M0491S). Vectors were digested using enzymatic restriction digest and ligated to gel purified PCR products using Gibson assembly. Ligated plasmid products were transformed into competent XL1-Blue *E. coli*. Detailed amino acid sequences are given in the Supplementary Information.

### Mammalian cell culture and transfection

HEK293T cells from ATCC (<30 passages) were cultured as a monolayer in complete media (Dulbecco’s modified Eagle’s medium (DMEM, Gibco Cat# 11965-092) supplemented with 10% (w/v) fetal bovine serum (FBS, VWR Cat# 97068-085) and 1% penicillin-streptomycin (VWR Cat# 16777-164)) at 37 ℃ under 5% CO_2_. In all experiments, glass coverslips and plates were pre-treated with 10 ug/mL human fibronectin (Millipore Cat# FC010) in Dulbecco’s PBS (DPBS, Gibco Cat# 14190-144) for 30 min at room temperature before cell plating. For transient expression, cells were transfected at 50-70% confluency with indicated expression plasmids using PEI (Polysciences Cat# 24765-1) in DMEM without FBS.

### Generation of VAMP2-expessing reporter HEK293T cells

Reporter cells expressing mCherry-myc-VAMP2 were generated by lentiviral transduction and antibiotic selection. For lentivirus generation, HEK293T cells were cultured in 6-well plates and were transfected at approximately 70% confluency with 1000 ng of the lentiviral vector of interest, packaging plasmids pCMV-dR8.91 (900 ng), and pCMV-VSV-G (100 ng) with 12 µL of PEI. Approximately 48 h after transfection, the cell medium was collected, filtered through a 0.45 µm filter, and then stored in 200 µL aliquots (“supernatant lentivirus”). The aliquots were flash-frozen in liquid nitrogen and stored at -80 ℃. For lentiviral transduction, HEK293T cells (<10 passages) were cultured in 6-well plates and transduced with 200 µL of supernatant lentivirus at approximately 50% confluency. Two days after transduction, cells with stably integrated reporters were selected by maintaining the cells in complete media containing antibiotics (250 µg/mL hygromycin for mCherry-myc-VAMP2). Antibiotic selection was performed until all cells showed mCherry signal under a tabletop fluorescence microscope. After selection, reporter cell lines were grown in T25 tissue culture flask until confluency, and frozen stocks were prepared and stored in liquid nitrogen for future use.

### Rat cortical neuron culture and AAV infection

All procedures were approved and carried out in compliance with the Stanford University Administrative Panel on Laboratory Animal Care, and all experiments were performed in accordance with relevant guidelines and regulations (protocol no. APLAC-32980). Before plating, plates were coated with 0.001% (w/v) poly-L-ornithine (Sigma-Aldrich Cat# P4957) in DPBS at room temperature overnight, washed twice with DPBS, and subsequently coated with 3.3 µg/mL of mouse laminin (Gibco Cat# 23017015) in DPBS at 37 ℃ for 2-4 h. After laminin coating, coverslips were washed twice with DPBS and stored at 4 ℃ until plating. Cortical neurons were extracted from embryonic day 18 Sprague Dawley rat embryos (Charles River Laboratories) by dissociation in Hank’s balanced salt solution (HBSS, Gibco Cat# 14025076). Cortical tissue was digested in papain according to the manufacturer’s protocol (Worthington Cat# LK003150), then plated onto 0.1 mm thick glass coverslips in complete neurobasal medium (CNB) at 37 ℃ under 5% CO2. CNB is neurobasal (Gibco Cat# 21103049), supplemented with 2% (v/v) B27 supplement (Life Technologies Cat# A3582801), 0.1% (v/v) FBS, 1% GlutaMAX (Gibco Cat# 35050061), 1% penicillin-streptomycin, and 1 mM sodium pyruvate (Gibco Cat# 11360070). On DIV1, half of the media was removed from each well and replaced with equal volume of CNB. On DIV4, neurons were infected with purified AAVs along with a media change. Neurons were wrapped in aluminum foil and were allowed to express for an additional 7 days in the incubator until light stimulation and subsequent analysis.

### Light stimulation of LATeNT in cultured mammalian cells and neurons

After cells were transfected with LATeNT plasmids or transduced with LATeNT AAVs, the plates were covered with aluminum foil until ready for blue light stimulation. When uncovered, cells were handled in a dark room under red light to avoid undesired activation of LATeNT. To stimulate LATeNT, cells were incubated at 37 ℃ while being placed directly on top of a blue light LED array. We used an AMUZA system consisting of a blue LED array, an LED Array Driver and pulse generator. We used a 16% duty cycle (2 sec on and 10 sec off) for 30 min, unless noted otherwise. Light power before pulse generation was typically around 1.0 mW/cm^2^ (measured by Coherent Fieldmax^TM^ II TO laser power meter), unless noted otherwise. Control samples were processed in parallel omitting light. Following light stimulation, cells were once again handled under red light, washed with DPBS three times and analyzed by western blot or immunofluorescence as described below.

### Western blot detection of LATeNT activity using anti-cleaved VAMP2 antibody

Following light stimulation and DPBS washes, cells were lysed directly in the cell culture wells with RIPA lysis buffer supplemented with 1x protease inhibitor cocktail (PIC, Thermo Scientific Cat# 78429) and 10 mM of picolinic acid (Sigma-Aldrich). Lysates were cleared via centrifugation at 20,000g at 4 ℃ for 10 min. Cleared lysates were mixed with protein loading buffer and boiled at 95 ℃ for 10 min. The concentrations of cell lysates were normalized using a Pierce^TM^ BCA Protein Assay Kit (Pierce Cat# 23225) and samples were loaded onto a polyacrylamide gel, and transferred onto PVDF membranes (Cytiva Cat# 10600023). Blots were blocked in 2% (w/v) nonfat milk (Lab Scientific, M-0841) in 1× tris-buffered saline with Tween (TBST) (Teknova Cat# T1688PK) for 15 min at room temperature, incubated with mouse antibody recognizing the N-terminal fragment of cleaved VAMP2 (VAMP/B/1148)^61^ for overnight at 4 ℃ in the blocking buffer. After primary antibody staining, the membrane was washed three times with TBST for 5 min each, incubated in anti-mouse-HRP secondary antibody for 90 min at room temperature in 2% (w/v) bovine serum albumin (BSA, Fischer Scientific Cat# BP1600-1) in TBST. After secondary antibody staining, the membrane was washed three times with TBST, rinsed twice with dH2O, and chemiluminescence was developed with Clarity^TM^ Western ECL Substrate (Bio-Rad) and imaged using a ChemiDoc XRS (Bio-Rad) imaging system. Except for the VAMP/B/1148 antibody, other primary antibody stainings, including the antibody that recognizes the C-terminal fragment of cleaved VAMP2 (VAMP/B/151)^61^, were done in 2% BSA in TBST, either for 2 h at room temperature or for overnight at 4 ℃.

### Immunofluorescence detection of LATeNT activity using anti-cleaved VAMP2 antibody

For confocal fluorescence microscopy experiments, cells were grown on 12mm-diameter glass coverslips (VWR Cat# 72230-01) in 24-well plates. Cells were transfected with LATeNT expression plasmids and stimulated with blue light as described above. After light stimulation, cells were gently washed three times with DPBS and were fixed with 4% (v/v) paraformaldehyde (PFA, ChemCruz Cat# sc-281692) at room temperature for 15 min. After fixation, PFA was aspirated and cells were permeabilized for 5 min with ice-cold methanol. Cells were washed again three times with DPBS and blocked for 1 h with 1% bovine serum albumin (BSA, Fischer Scientific Cat# BP1600-1) in TBST at 4 ℃. Cells were then incubated with primary antibodies in TBST overnight at 4 °C. In immunofluorescence detection of LATeNT activity, mouse antibody recognizing the C-terminal fragment of cleaved VAMP2 (VAMP/B/151)^61^ was used, as VAMP/B/1148 staining was not detectable in fixed cells. After washing three times with TBST, cells were incubated with fluorophore-conjugated secondary antibodies in TBST for 1 h at room temperature. Cells were washed three times with TBST and imaged.

Imaging was performed with a Zeiss Axio Observer.Z1 microscope with a Yokogawa spinning disk confocal head, Cascade IIL:512 camera, a Quad-band notch dichroic mirror (405/488/568/647 nm), and 405 nm, 491 nm, 561 nm, and 640 nm lasers (all 50 mW). Images were captured through a 63× oil-immersion objective for the following fluorophores: AlexaFluor 405 (405 laser excitation, 445/40 emission), EGFR and AlexaFluor 488 (491 laser excitation, 528/38 emission), mCherry and AlexaFluor 568 (561 laser excitation, 617/73 emission), and AlexaFluor 647 (647 laser excitation, 700/75 emission). Image acquisition times ranged from 10 to 500 ms per channel, and images were captured as the average of two or three such exposures in rapid succession. Image acquisition and processing was carried out with the SlideBook 5.0 software (Intelligent Imaging Innovations, 3i).

### Construction of yeast strains

All strains were derived from *Saccharomyces cerevisiae* strain BY4741. Plasmid transformation or integration in yeast was performed using the Frozen E-Z Yeast Transformation II kit (Zymo Research Cat# T2001) according to the manufacturer’s protocol. *S. cerevisiae* strains were produced stepwise and propagated at 30 ℃ in complete minimal media (CSM) with 20g/L dextrose (CSM-D). CSM is 6.7 g/L yeast nitrogen base without amino acids (Thermo Fischer Scientific, Cat# H26271) and 0.54 g/L CSM-Ade-His-Leu-Lys-Trp-Ura (Sunrise Science Products Cat# 1135-010).

Transformants were isolated in appropriate selective medium by auxotrophic complementation. For yeast strain transformation, we grew cells at 30 ℃ in YPD containing 10 g/L yeast extract (Gibco Cat# 212750), 20 g/L peptone (Gibco Cat# 211677), and 20 g/L dextrose. We first obtained the yeast strain containing the membrane tethered transcription factor and the reporter gene, by integrating the plasmid containing lexO::YFP-URA and ACT1::LexA-VP16- VAMP2, along with a HIS3 gene. Transformed cells containing the desired integration were selected on CSM plates supplemented with 100 mg/L leucine and 800 mg/L uracil (CSM-D + Leu, Ura) With lexO::YFP reporter cells in hand, we episomally introduced plasmids containing GAL1::TeNT-mCherry or GAL1::LATeNT1-mCherry, along with a LEU2 gene. Transformed yeast cells containing the plasmid were selected on CSM plates supplemented with 800 mg/L uracil.

### Yeast culture and Analysis of LATeNT activity in yeast cytosol

Yeast strains containing the YFP reporter, VAMP2-tethered transcription factor, and galactose-inducible LATeNT-mCherry (or TeNT-mCherry) were propagated at 30 ℃ in CSM-D media supplemented with 800 mg/L uracil (CSM-D + Ura). To induce protein expression, yeasts were inoculated from saturated cultures in CSM with 2 g/L dextrose and 18 g/L galactose (CSM- D/G) overnight from a 1:20 dilution. After inoculation, culture tubes were wrapped in aluminum foil to prevent light exposure.

After overnight induction, the culture was exposed to ambient room light for 1-10 min, wrapped in aluminum foil, and cultured at 30 ℃ for additional 8 h to allow reporter expression. Cells were then collected from 1 mL of cell culture, by pelleting at 3,000g for 2 min at 4 ℃ and resuspending in 0.1 ml PBS with 0.1% BSA (PBS-B). To analyze and sort, single yeast cells were gated on a forward-scatter area (FSC-A) by side-scatter area (SSC-A) plot around the clustered population (P1) on a ZE5 Cell Analyzer (Bio-Rad). P1 was then gated on a FSC-A by forward-scatter height (FSC-H) around the clustered population (P2). P2 populated cells were then plotted to detect mCherry and YFP signals. FlowJo v10 (BD Biosciences) was used to analyze FACS data.

### Library generation for directed evolution of LATeNT in yeast

Mutant LATeNT1 library was generated using error-prone PCR with 100 ng of plasmid encoding GAL1::mCherry-LATeNT1 as the template. Three libraries were generated with different levels of mutagenesis, by carrying out error-prone PCR with different levels of mutagenesis:

E1-1 (low mutation): 5 μM 8-oxo-dGTP, 1 μM dPTP, 15 PCR cycles

E1-2 (medium mutation): 10 μM 8-oxo-dGTP, 2 μM dPTP, 15 PCR cycles E1-3 (high mutation): 15 μM 8-oxo-dGTP, 4 μM dPTP, 15 PCR cycles

Error-prone PCR was carried out following published protocols^62^, with following primers that restricted the mutation to the LATeNT1 region:

F: GCCACCatgGCTAGCGTTAACccgatcacc

R: ttgtcctcctcgcccttgctca

PCR products were gel purified then reamplified for 30 more cycles under normal conditions and gel purified again. The original plasmid was digested with NheI and MscI and gel purified as well, to serve as a backbone for DNA recombination. Both 4,000 ng of PCR product and 1,000 ng of cut vector were mixed and water was added to total 10 μL. The resulting mixture was electroporated into electrocompetent lexO::YFP reporter yeasts. After electroporation, cells were rescued in 2 mL of YPD media and recovered at 30 ℃ for 1 h. Then, 1.98 mL of the culture was propagated in 100 mL of CSM-D +Ura media, while the remaining 20 µl was used to determine library size. Yeasts were diluted 100×, 1,000×, 10,000×, 100,000× and 20 µl of each dilution was plated onto CSM-D + Ura plates at 30 ℃ for 3 days. The resulting library size was determined by the number of colonies on the 100×, 100×, 1,000× and 100,000× plates, corresponding to 10^4^, 10^5^, 10^6^, or 10^7^ transformants in the library, respectively. The library sizes resulting from the three libraries were 3.5 x 10^7^, 4.0 x 10^7^, and 4.0 x 10^7^, respectively. All three libraries were tested and showed expected light-dependent YFP expression compared to the original LATeNT1 strain. All libraries were combined with equal cell numbers to perform subsequent directed evolution.

### Directed evolution of LATeNT in yeast

Yeast cells were sorted using Sony SH800 sorter (Sony Biotechnology). Gates for singlets were drawn using FSC-A, SSC-A, and FSC-H values as previously described. The resulting singlet population was drawn on a mCherry by YFP plot to collect cells with desired activity and expression level.

In round 1, cells were induced while exposed to ambient light. We collected the top 1.03% of cells with high YFP/mCherry ratio, as these cells will contain LATeNT variant that is active under light.

In round 2 to 5, cells were induced while wrapped in aluminum foil. Two different selections were done alternatively, either sorting cells with high mCherry/YFP ratio after light stimulation (positive control), or sorting cells with low YFP/mCherry ratio without light stimulation (negative control). Each selection was performed to enrich LATeNT mutant with high light-dependent activity, or with low background activity at dark. Round 2 and 4 were negative selection with top 2.2% and 2.6% of cells being collected, respectively. Round 3 and 5 were positive selection with top 1.7% and 0.4% of cells being collected, respectively.

After 5 rounds of enrichment, plasmids in the enriched library was extracted using the Zymoprep yeast Plasmid Miniprep II kit (Zymo Research Cat# D2004) following manufacturer’s protocol. Then, plasmids were transformed into competent XL1-Blue *E. coli*, single colonies were grown overnight, and plasmid was extracted and sequenced by Sanger sequencing.

### *In vitro* VAMP2 cleavage assay by TeNT and LATeNT

For in vitro VAMP2 cleavage assay, wild-type HEK293T cells were transfected with mCherry- myc-VAMP2, TeNT, or LATeNT expression plasmid, as described above, and cells were kept in the dark. 24 h after transfection, cells were stimulated with light for 30 min and cytosolic proteins were extracted using M-PER^TM^ mammalian protein extraction reagent (Thermo Scientific, Cat# 78501) under ambient room light. In some cases, proteins were extracted in the dark without light stimulation (see **Fig. S3B**). The soluble fraction of VAMP2 lysate and TeNT (or LATeNT) lysate were mixed and incubated under ambient room light. The reaction was quenched by adding 50 mM of picolinic acid to inhibit protease activity. The reaction mixture was further analyzed by Western blotting.

### BRET activation of NanoLuc-LATeNT, Ca-LATeNT, or GPCR-LATeNT*

For LATeNT activation by BRET, cells were cultured and transfected with LATeNT constructs as described above. 24 h after transfection, the media was replaced with a media containing 50 μM of fluorofurimazine (FFz, Selleck Chemicals Cat# E1620) in the dark room and incubated at 37 ℃ for 1 h. For Ca^2+^ stimulation, CaCl_2_ and ionomycin were added to a final concentration of 5 mM and 2 μM, respectively. For GPCR activation, GPCR agonists CCL20 or Salvinorin B was added to a final concentratin of 0.2 μg/mL or 1 μM, respectively. After FFz treatment, cells were washed with DPBS and analyzed by Western blotting as described above.

### Light-dependent reporter expression using Gal4-LATeNT*

For Gal4-LATeNT* experiments, HEK293T cells were cultured on coverslips and transfected with Gal4-VAMP2-LATeNT* constructs as described above. 24 h after transfection, cells were stimulated with light for 30 min and kept in dark for additional 24 h to allow reporter expression. Cells were PFA-fixed in the dark room and subsequently stained with anti-HA antibody to visualize reporter expression. mCherry expression was quantified by confocal imaging.

### RELEASE-LATeNT assay

For RELEASE-LATeNT, cells were cultured, transfected with LATeNT constructs, and stimulated with light as described above. After light stimulation, cells were covered with aluminum foil and incubated at 37 ℃ for 4 h. SEAP activity in the cell culture supernatant was measured following published protocols^50^.

### Glucose-stimulated insulin secretion (GSIS) assay

MIN6 cell culture and GSIS assay were performed following published protocols^45, 63^. Briefly, cells were maintained in DMEM supplemented with 15% heat-inactivated FBS, 2 mM GlutaMAX, 1 mM sodium pyruvate, 1 % penicillin-streptomycin, and 0.05 mM beta-mercaptoethanol. MIN6 cells were transfected with doxycycline-inducible LATeNT using lentiviral delivery and selected by puromycin. For GSIS assay, cells were starved for 1 hour under KRB buffer^63^ containing 1 mM glucose. After starvation, media was replaced with KRB buffer with either low (1 mM) or high (25 mM) glucose concentration. After 1 hour, cell culture supernatant was collected, and remaining cells were lysed. The amount of insulin in the supernatant and the lysate was measured using insulin ELISA kit (Mercodia) following manufacturer’s protocols. For LATeNT-expressing MIN6 cells, cells were either kept in dark or stimulated for 30 min with ambient room light prior to starvation.

### Animals

*Sst*-IRES-FlpO (028579, Jackson Research laboratories), *Sst*-IRES-Cre (013044, Jackson Research Laboratories) and C57BL/6J mice (purchased from Daehan Biolink) used in the current study were maintained and handled in accordance with protocols (DGIST-IACUC- 20122401-0004) approved by the Institutional Animal Care and Use Committee (IACUC) of the Daegu Gyeongbuk Institute of Science and Technology (DGIST). Mice were maintained on a 12:12 h light:dark cycle under standard temperature (22 ± 2 ℃)-controlled laboratory conditions and received water and food ad libitum. Pregnant rats purchased from Daehan Biolink were used for *in vitro* culture of dissociated hippocampal neurons. All experimental procedures were conducted according to guidelines and protocols for rodent experimentation approved by the IACUC of DGIST.

### Electrophysiology recordings in cultured hippocampal neurons

Cultured hippocampal neurons were infected at DIV4 with AAVs expressing mCherry-LATeNT and kept in the dark. At DIV14–16, neurons were either kept in the dark or stimulated with 470 nm LED light for 30 min and analyzed using whole-cell patch-clamp electrophysiological recordings. Pipettes were pulled from borosilicate glass (o.d. 1.5 mm, i.d. 0.86 mm; Sutter Instruments) using a Model P-97 pipette puller (Sutter Instruments). The resistance of patch pipettes filled with internal solution varied between 3 and 6 MΩ. For recordings of mEPSCs, the composition of the internal solution was 145 mM CsCl, 5 mM NaCl, 10 mM HEPES, 10 mM EGTA, 0.3 mM Na-GTP, and 4 mM Mg-ATP, with pH adjusted to 7.2–7.4 with CsOH and an osmolarity of 290–295 mOsmol/L. The external solution consisted of 130 mM NaCl, 4 mM KCl, 2 mM CaCl_2_, 1 mM MgCl_2_, 10 mM HEPES, and 10 mM D-glucose, with pH adjusted to 7.2–7.4 with NaOH and an osmolarity of 300–305 mOsmol/L. The whole-cell configuration was obtained at room temperature using μM-TSC manipulators (Sensapex). Electrophysiological data were acquired with a Multiclamp 700B amplifier (Axon Instruments) and pCLAMP software and digitized using an Axon DigiData 1550B data acquisition board (Axon Instruments). mEPSCs were recorded at a holding potential of -70 mV. Synaptic currents were analyzed offline using Clampfit 10.8 software (Molecular Devices). The external solution contained 1 μM TTX and 50 μM picrotoxin to block GABA_A_ receptor and Na^+^ currents, respectively.

### Electrophysiology recordings in acute brain slices

Six-wk-old C57BL/6J or *Sst*-IRES-FlpO mice were injected in the hippocampal CA3 region with AAV-mCherry LATeNT, and transverse hippocampal slices were prepared 2 wk later. After anesthesia with isoflurane, mice were decapitated and their brains were rapidly removed and placed in an ice-cold oxygenated (95% O_2_ and 5% CO_2_), low-Ca^2+^/high-Mg^2+^ solution (3.3 mM KCl, 1.3 mM NaH_2_PO_4_, 26 mM NaHCO_3_, 11 mM D-glucose, 211 mM sucrose, 0.5 mM CaCl_2_ and 10 mM MgCl_2_). Hippocampal slices were cut using a vibratome (WT1000s, Leica) and transferred to a holding chamber containing oxygenated artificial cerebrospinal fluid (aCSF; 124 mM NaCl, 3.3 mM KCl, 1.3 mM NaH_2_PO_4_, 26 mM NaHCO_3_, 11 mM D-glucose, 3 mM CaCl_2_ and 1.5 mM MgCl_2_). For the control group, slices were incubated at 30 ℃ for at least 60 min and used for experiments within 4 h under dark conditions. For the experimental group in **Fig. 3A-F**, slices were stimulated under ambient room light (∼0.225 mW/cm^2^) for 1-2 hours before recordings. For the experimental groups in **Fig. 3G-H**, samples were maintained in darkness for a 5-min baseline recording and then either kept in darkness or exposed to ambient light. All experiments were performed at 30–32 ℃. Recordings were performed using a Multiclamp 700B amplifier and DigiData 1550B digitizer (Molecular Devices). For measurement of evoked EPSCs (eEPSCs), slices were placed in the recording chamber and perfused continuously with 95% O_2_/5% CO_2_-bubbled aCSF. Patch pipettes (3–5MΩ) were filled with an internal solution consisting of 130 mM Cs-methanesulfonate, 5 mM TEA-Cl, 8 mM NaCl, 0.5 mM EGTA, 10 mM HEPES, 4 mM Mg-ATP, 0.4 mM Na-GTP, 1 mM QX-314, and 10 mM disodium phosphocreatine. The osmolarity of the internal solution was 280–290 mOsm. Electrical stimulation was applied using a concentric bipolar electrode (FHC), placed in the striatum radiatum. eEPSCs were recorded at -70 mV (for AMPA-EPSCs) and +40 mV (for NMDA-EPSCs; 50 ms after stimulation). Input/output (I/O) responses were obtained by eliciting EPSCs with a series of stimulation intensities (20-100 μA). Paired-pulse ratios (PPRs) were calculated by delivering two pulses at different intervals (50, 100, 200, and 500 ms). For eEPSC recordings, PTX (50 μM) was applied to block the GABA_A_ receptor. Only cells with an access resistance (Ra) in the range of 5 MΩ < Ra < 30 MΩ were analyzed. To measure changes in eEPSC kinetics, a stable baseline was obtained under the dark condition for 5 min, and then 473-nm blue light (0.5 mW/cm²) was applied for 30 min. For measurement of evoked IPSCs (eIPSCs), slices were placed in the recording chamber and perfused continuously with 95% O_2_/5% CO_2_-bubbled aCSF. Patch pipettes (3–8MΩ) were filled with an internal solution consisting of 130 mM Cs-methanesulfonate, 5 mM TEA-Cl, 8 mM NaCl, 0.5 mM EGTA, 10 mM HEPES, 4 mM Mg-ATP, 0.4 mM Na-GTP, 1 mM QX-314, and 10 mM disodium phosphocreatine. Electrical stimulation was applied using a concentric bipolar electrode (FHC), placed in the stratum pyramidale, or stratum lacunosum moleculare. eIPSCs were recorded at 0 mV. Input/output (I/O) responses were obtained by eliciting IPSCs with a series of stimulation intensities (10-60 μA). Paired-pulse ratios (PPRs) were calculated by delivering two pulses at different intervals (50, 100, 200, and 500 ms). For IPSC recordings, D-AP5 (50 μM) and CNQX (10 μM) were applied to block the NMDA and AMPA receptors respectively. Only cells with an access resistance (Ra) in the range of 5 MΩ < Ra < 30 MΩ were analyzed. For measurement of neurotransmission in long-range projections, mice were injected in the vHPP (AP, -3.5 mm; ML, ±3.5 mm; DV, -3.8 mm) with AAV-mCherry LATeNT and maintained for 3 weeks, and transverse hippocampal slices were prepared. To obtain coronal slices of the mPFC containing the long projection from the vHPP to the mPFC, each whole brain was cut at a 10° angle relative to the coronal plane, beginning at the front of the forebrain. mEPSC recordings were performed as described above. For eNpHR validation experiments, 300 pA current injections were delivered into CA1 stratum oriens SST+ interneurons and action potential firing was recorded. Electrophysiological data were acquired using the pCLAMP software and a MultiClamp 700B (Axon Instruments), and were digitized using an Axon DigiData 1550B data acquisition board (Axon Instruments). Data were sampled at 10 kHz and filtered at 4 kHz. Data were discarded if the series resistance was greater than 30 MΩ or differed by more than 20%.

### Preparation and titration of adeno-associated viruses

AAVs were prepared as described previously^64^. In brief, HEK293T cells were co-transfected with pHelper and pAAV1.0, together with pAAV-CAG-mCherry-LATeNT or pAAV-EF1a-fDIO-mCherry-LATeNT. Cells were harvested 72 h later, lysed, mixed with 40% polyethylene glycol and 2.5 M NaCl, and centrifuged at 2,000 × g for 10 min. The resulting pellets were resuspended in HEPES buffer (20 mM HEPES, 115 mM NaCl, 1.2 mM CaCl_2_, 1.2 mM MgCl_2_, 2.4 mM KH_2_PO_4_), mixed with an equal volume of chloroform, and centrifuged at 400 × g for 7 min. The supernatant fractions were concentrated with a Centriprep centrifugal filter (Millipore 4310) at 2,500 rpm for 15 min each, followed by concentration with an Amicon Ultra centrifugal filter (0.5 mL) at 14,000 rpm for 10 min. Virus infection titers were determined by qRT-PCR detection of mCherry sequences based on a standard curve generated using the corresponding DNA plasmid.

### Stereotaxic surgery and virus injections

Six-wk-old C57BL/6J, *Sst*-IRES-FlpO or *Sst*-IRES-Cre mice, with heads firmly secured in a stereotactic device, were anesthetized by inhalation of isoflurane (2–3%). Virus solutions were injected using a Hamilton syringe at a flow rate of 0.1 µL/min. The indicated coordinates were used for stereotaxic injection into the hippocampal CA1 (AP, -2.5 mm; ML, ±1.5 mm, DV, -1.3 mm) or hippocampal CA3 (AP, -2.1 mm; ML, ±2.3 mm; and DV, -2.4 mm). After 2-wk recovery period, the injected regions were subjected to *ex vivo* electrophysiological analyses. For *in vivo* experiments, optical fibers (FC-ZF1.25F; Doric Lenses Inc.) were implanted perpendicularly into the targeted brain region 1 wk after viral injection. Mice with off-target viral infection, identified by *post hoc* histological analysis, were excluded from all analyses.

### Enriched environmental protocol

Three weeks after viral injection, blue light (473 nm, 1 mW) was delivered to the dSub region for 30 min (2 sec on/10 sec off), after which mice were exposed to EE conditions for 8 h per day for 5 consecutive days, then subjected to functional analyses. Blue laser (BL473T3-100FC; Shanghai Laser & Optics Century) was controlled with an isolated pulse stimulator (MODEL 2100; A-M SYSTEMS) connected to the optic fiber (FC-ZF1.25F; Doric Lenses Inc). The EE consisted of a large cage containing a running wheel, hut, tunnel, and several other novel objects, as previously described^37^.

### Elevated plus maze test

The elevated plus-maze is a plus-shaped (+), white, acrylic maze with two open arms (30 × 5 × 0.5 cm) and two closed arms (30 × 5 × 30 cm) positioned at a height of 75 cm from the floor. Light conditions around open and closed arms were ∼300 and ∼30 lux, respectively. For the test, mice were introduced into the center zone of the elevated plus-maze and allowed to move freely for 5 minutes. For LATeNT-mediated inhibition, blue light (473 nm, 1 mW) was delivered to the hippocampal CA1 region for 30 min (2 sec on/10 sec off) before mice were tested in the EPM. For NpHR-mediated inhibition, continuous yellow light (594 nm, 5 mW) was delivered to the hippocampal CA1 region for 5 min while mice were tested in the EPM. All behaviors were recorded by a top-view infrared camera, and the time spent in each arm was measured and analyzed using EthoVision XT 10.5 software (Noldus).

### Bioluminescence imaging of mice

Three weeks after injecting AAV-NanoLuc-LATeNT* into the hippocampal CA3 regions, mice were administered intraperitoneal (i.p.) injections of reconstituted CFz (1.3 µM), as previously described^43^. Bioluminescence images were acquired using the IVIS Spectrum In Vivo Imaging System (Caliper Life Sciences). During imaging, mice were anesthetized with isoflurane delivered via an XGI-8 Gas Anesthesia System (Caliper Life Sciences). Image settings were as follows: emission filter open, field of view of 14 cm, f-stop of 1.0, height of 2.5 cm, and exposure time of 1 s.

### Immunohistochemistry and imaging

Mice were anesthetized and immediately perfused, first with PBS for 3 minutes and then with 4% paraformaldehyde for 5 minutes. Brains were dissected out, fixed in 4% paraformaldehyde overnight, and then incubated with 30% sucrose (in PBS) overnight, and sliced into 40 μm- thick coronal sections using a vibratome (Model VT1200S; Leica Biosystems). Sections were permeabilized by incubating with 0.2% Triton X-100 in PBS containing 5% bovine serum albumin and 5% horse serum for 1 hour. For immunostaining, sections were incubated for 8– 12 hours at 4℃ with primary antibodies diluted in the same blocking solution. The following primary antibodies were used: anti-c-Fos (Cell signaling Cat#2250, RRID: AB_2247211, 1: 500), or anti-SST (Millipore Cat# MAB354, RRID: AB_2255365, 1:50) antibodies. Sections were washed three times in PBS and incubated with appropriate Cy3- or FITC-conjugated secondary antibodies (Jackson ImmunoResearch) for 2 hours at room temperature. After three washes with PBS, sections were mounted onto glass slides (Superfrost Plus; Fisher Scientific) with Vectashield mounting medium (H-1200; Vector Laboratories). Z stack images (5 images, 3 μm thickness) were acquired in standard mode with a laser-scanning confocal microscope (LSM800 with Airyscan mode; Zeiss) and processed using the maximum intensity projection function in Zen2.6 software (Zeiss).

### Statistical analyses

Data were assessed by one-way analysis of variance (ANOVA) with Tukey’s *post hoc* comparisons or Mann–Whitney *U* test, or Friedman test; ‘*n*’ numbers and tests used to determine statistical significance were stated in the figure legends. The normality of data distributions was evaluated using the Shapiro-Wilk test. Prism 10 (GraphPad Software) was used for the analysis of data and preparation of bar graphs. p-values < 0.05 were considered statistically significant.

## References

1. Rizo, J. Molecular Mechanisms Underlying Neurotransmitter Release. Annu Rev Biophys 51, 377–408 (2022).

2. Dong, C. Cytokine Regulation and Function in T Cells. Annu Rev Immunol 39, 51–76 (2021).

3. Dong, M., Masuyer, G. &Stenmark, P. Botulinum and Tetanus Neurotoxins. Annu Rev Biochem 88, 811–837 (2019).

4. Réthy, L. & Réthy, L.A. Human lethal dose of tetanus toxin. Lancet 350, 1518 (1997).

5. Humeau, Y., Doussau, F., Grant, N.J. & Poulain, B. How botulinum and tetanus neurotoxins block neurotransmitter release. Biochimie 82, 427–446 (2000).

6. Cornille, F. et al. Cooperative exosite-dependent cleavage of synaptobrevin by tetanus toxin light chain. J Biol Chem 272, 3459–3464 (1997).

7. Sikorra, S., Henke, T., Galli, T. & Binz, T. Substrate recognition mechanism of VAMP/synaptobrevin-cleaving clostridial neurotoxins. J Biol Chem 283, 21145–21152 (2008).

8. Chen, S., Hall, C. & Barbieri, J.T. Substrate recognition of VAMP-2 by botulinum neurotoxin B and tetanus neurotoxin. J Biol Chem 283, 21153–21159 (2008).

9. Schiavo, G., Rossetto, O., Benfenati, F., Poulain, B. & Montecucco, C. Tetanus and botulinum neurotoxins are zinc proteases specific for components of the neuroexocytosis apparatus. Ann N Y Acad Sci 710, 65–75 (1994).

10. Nakashiba, T., Young, J.Z., McHugh, T.J., Buhl, D.L. & Tonegawa, S. Transgenic inhibition of synaptic transmission reveals role of CA3 output in hippocampal learning. Science 319, 1260–1264 (2008).

11. Macosko, E.Z. et al. A hub-and-spoke circuit drives pheromone attraction and social behaviour in C. elegans. Nature 458, 1171–1175 (2009).

12. Miyashita, S.I., Zhang, J., Zhang, S., Shoemaker, C.B. & Dong, M. Delivery of single-domain antibodies into neurons using a chimeric toxin-based platform is therapeutic in mouse models of botulism. Sci Transl Med 13 (2021).

13. McNutt, P.M. et al. Neuronal delivery of antibodies has therapeutic effects in animal models of botulism. Sci Transl Med 13 (2021).

14. Tian, S. et al. Targeted intracellular delivery of Cas13 and Cas9 nucleases using bacterial toxin-based platforms. Cell Rep 38, 110476 (2022).

15. Roh, H., Dorner, B.G. & Ting, A.Y. Cell-Type-Specific Intracellular Protein Delivery with Inactivated Botulinum Neurotoxin. J Am Chem Soc 145, 10220–10226 (2023).

16. Francis, J.W., Hosler, B.A., Brown, R.H. & Fishman, P.S. CuZn superoxide dismutase (SOD- 1):tetanus toxin fragment C hybrid protein for targeted delivery of SOD-1 to neuronal cells. J Biol Chem 270, 15434–15442 (1995).

17. Blum, T.R. et al. Phage-assisted evolution of botulinum neurotoxin proteases with reprogrammed specificity. Science 371, 803–810 (2021).

18. Regazzi, R. et al. VAMP-2 and cellubrevin are expressed in pancreatic beta-cells and are essential for Ca(2+)-but not for GTP gamma S-induced insulin secretion. EMBO J 14, 2723–2730 (1995).

19. McCue, A.C. & Kuhlman, B. Design and engineering of light-sensitive protein switches. Curr Opin Struct Biol 74, 102377 (2022).

20. Cho, K.F. et al. Split-TurboID enables contact-dependent proximity labeling in cells. Proc Natl Acad Sci U S A 117, 12143–12154 (2020).

21. Lim, S.A. & Wells, J.A. Split enzymes: Design principles and strategy. Methods Enzymol 644, 275–296 (2020).

22. Sanchez, M.I., Nguyen, Q.A., Wang, W., Soltesz, I. & Ting, A.Y. Transcriptional readout of neuronal activity via an engineered Ca. Proc Natl Acad Sci U S A 117, 33186–33196 (2020).

23. Lee, S.Y. et al. Engineered allostery in light-regulated LOV-Turbo enables precise spatiotemporal control of proximity labeling in living cells. Nat Methods 20, 908–917 (2023).

24. Huala, E. et al. Arabidopsis NPH1: a protein kinase with a putative redox-sensing domain. Science 278, 2120–2123 (1997).

25. Swartz, T.E. et al. The photocycle of a flavin-binding domain of the blue light photoreceptor phototropin. J Biol Chem 276, 36493–36500 (2001).

26. Breidenbach, M.A. & Brunger, A.T. 2.3 A crystal structure of tetanus neurotoxin light chain. Biochemistry 44, 7450–7457 (2005).

27. Kim, M.W. et al. Time-gated detection of protein-protein interactions with transcriptional readout. Elife 6 (2017).

28. Gradinaru, V., Thompson, K.R. & Deisseroth, K. eNpHR: a Natronomonas halorhodopsin enhanced for optogenetic applications. Brain Cell Biol 36, 129–139 (2008).

29. Copits, B.A. et al. A photoswitchable GPCR-based opsin for presynaptic inhibition. Neuron 109, 1791–1809.e1711 (2021).

30. Wang, W. et al. A light- and calcium-gated transcription factor for imaging and manipulating activated neurons. Nat Biotechnol 35, 864–871 (2017).

31. Kawano, F., Aono, Y., Suzuki, H. & Sato, M. Fluorescence imaging-based high-throughput screening of fast- and slow-cycling LOV proteins. PLoS One 8, e82693 (2013).

32. Hussain, S. & Davanger, S. Postsynaptic VAMP/Synaptobrevin Facilitates Differential Vesicle Trafficking of GluA1 and GluA2 AMPA Receptor Subunits. PLoS One 10, e0140868 (2015).

33. Wilhelm, B.G. et al. Composition of isolated synaptic boutons reveals the amounts of vesicle trafficking proteins. Science 344, 1023–1028 (2014).

34. Grote, E., Hao, J.C., Bennett, M.K. & Kelly, R.B. A targeting signal in VAMP regulating transport to synaptic vesicles. Cell 81, 581–589 (1995).

35. Phillips, M.L., Robinson, H.A. & Pozzo-Miller, L. Ventral hippocampal projections to the medial prefrontal cortex regulate social memory. Elife 8 (2019).

36. Nithianantharajah, J. & Hannan, A.J. Enriched environments, experience-dependent plasticity and disorders of the nervous system. Nat Rev Neurosci 7, 697–709 (2006).

37. Kim, S. et al. Npas4 regulates IQSEC3 expression in hippocampal somatostatin interneurons to mediate anxiety-like behavior. Cell Rep 36, 109417 (2021).

38. Klaric, T.S. et al. A reduction in Npas4 expression results in delayed neural differentiation of mouse embryonic stem cells. Stem Cell Res Ther 5, 64 (2014).

39. Miles, R., Tóth, K., Gulyás, A.I., Hájos, N. & Freund, T.F. Differences between somatic and dendritic inhibition in the hippocampus. Neuron 16, 815–823 (1996).

40. Gradinaru, V. et al. Molecular and cellular approaches for diversifying and extending optogenetics. Cell 141, 154–165 (2010).

41. Kim, C.K., Cho, K.F., Kim, M.W. & Ting, A.Y. Luciferase-LOV BRET enables versatile and specific transcriptional readout of cellular protein-protein interactions. Elife 8 (2019).

42. Lee, S.Y. et al. Engineered allostery in light-regulated LOV-Turbo enables precise spatiotemporal control of proximity labeling in living cells. bioRxiv (2023).

43. Su, Y. et al. An optimized bioluminescent substrate for non-invasive imaging in the brain. Nat Chem Biol 19, 731–739 (2023).

44. Grant, C.S. Insulinoma. Best Pract Res Clin Gastroenterol 19, 783–798 (2005).

45. Wu, C.T. et al. Discovery of ciliary G protein-coupled receptors regulating pancreatic islet insulin and glucagon secretion. Genes Dev 35, 1243–1255 (2021).

46. Fink, T. & Jerala, R. Designed protease-based signaling networks. Curr Opin Chem Biol 68, 102146 (2022).

47. 47. Lambert, G.G., et al. CaBLAM! A high-contrast bioluminescent Ca. *bioRxiv* (2023).

48. Cho, K.F. et al. A light-gated transcriptional recorder for detecting cell-cell contacts. Elife 11 (2022).

49. Vardy, E. et al. A New DREADD Facilitates the Multiplexed Chemogenetic Interrogation of Behavior. Neuron 86, 936–946 (2015).

50. Vlahos, A.E. et al. Protease-controlled secretion and display of intercellular signals. Nat Commun 13, 912 (2022).

51. Roth, B.L. DREADDs for Neuroscientists. Neuron 89, 683–694 (2016).

52. Wiegert, J.S., Mahn, M., Prigge, M., Printz, Y. & Yizhar, O. Silencing Neurons: Tools, Applications, and Experimental Constraints. Neuron 95, 504–529 (2017).

53. Saloman, J.L. et al. Gi-DREADD Expression in Peripheral Nerves Produces Ligand-Dependent Analgesia, as well as Ligand-Independent Functional Changes in Sensory Neurons. J Neurosci 36, 10769–10781 (2016).

54. Wang, C.L. et al. Activity-dependent development of callosal projections in the somatosensory cortex. J Neurosci 27, 11334–11342 (2007).

55. Liu, Q. et al. A Photoactivatable Botulinum Neurotoxin for Inducible Control of Neurotransmission. Neuron 101, 863–875.e866 (2019).

56. Li, X.L., Tei, R., Uematsu, M. & Baskin, J.M. Ultralow Background Membrane Editors for Spatiotemporal Control of Phosphatidic Acid Metabolism and Signaling. ACS Cent Sci 10, 543–554 (2024).

57. Reynolds, J.A., Vishweshwaraiah, Y.L., Chirasani, V.R., Pritchard, J.R. & Dokholyan, N.V. An engineered N-acyltransferase-LOV2 domain fusion protein enables light-inducible allosteric control of enzymatic activity. J Biol Chem 299, 103069 (2023).

58. Lacy, D.B. & Stevens, R.C. Sequence homology and structural analysis of the clostridial neurotoxins. J Mol Biol 291, 1091–1104 (1999).

59. Asadi, F. & Dhanvantari, S. Pathways of Glucagon Secretion and Trafficking in the Pancreatic Alpha Cell: Novel Pathways, Proteins, and Targets for Hyperglucagonemia. Front Endocrinol (Lausanne) 12, 726368 (2021).

60. Lee, H., Hong, Y.J., Baik, S., Hyeon, T. & Kim, D.H. Enzyme-Based Glucose Sensor: From Invasive to Wearable Device. Adv Healthc Mater 7, e1701150 (2018).

61. von Berg, L. et al. Functional detection of botulinum neurotoxin serotypes A to F by monoclonal neoepitope-specific antibodies and suspension array technology. Sci Rep 9, 5531 (2019).

62. Colby, D.W. et al. Engineering antibody affinity by yeast surface display. Methods Enzymol 388, 348–358 (2004).

63. Yang, L. & Chen, W. Insulin secretion assays in an engineered MIN6 cell line. MethodsX 10, 102029 (2023).

64. Kim, D. et al. IQSEC3 Deletion Impairs Fear Memory Through Upregulation of Ribosomal S6K1 Signaling in the Hippocampus. Biol Psychiatry 91, 821–831 (2022).

