## Supplementary Figures 1-7, Materials for "Light-activated tetanus neurotoxin for conditional proteolysis and inducible synaptic inhibition *in vivo*"

for

[††]Present address: Aperture Therapeutics, 733 Industrial Road, San Carlos, CA 94070, USA.

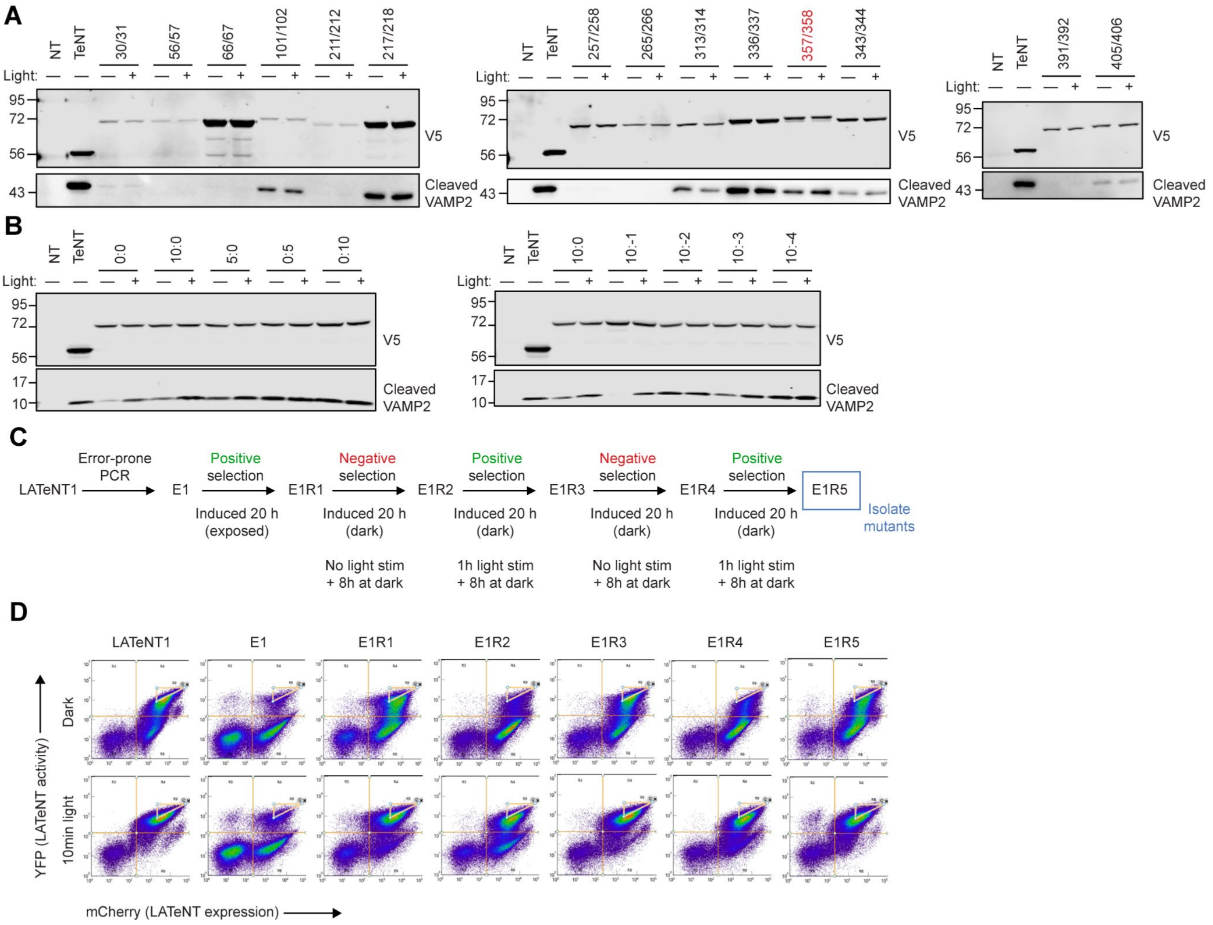


**Supplementary Figure 1. Design and directed evolution of LATeNT. Related to Figure 1.** **(A, B)** Western blots showing activities of (A) different LOV domain insertion constructs from Fig. 1B, C and (B) linker optimization constructs from Fig. 1D. VAMP2 reporter HEK293T cells expressing each construct were either kept in the dark or stimulated for 30 min with ambient room light. This experiment was performed once. **(C)** Directed evolution of LATeNT in yeast. LATeNT expression was induced for 20 h. During induction, cells were either exposed to room light (round 1) or were kept in the dark (rounds 2-5). In rounds 2-5, LATeNT-expressing cells were stimulated for 1 h with (rounds 3, 5) or kept in the dark (rounds 2, 4) following induction, and incubated for an additional 8 h in the dark before FACS enrichment. **(D)** FACS plots of enriched populations after each round of selection. Yeast cells were kept in the dark or stimulated for 10 min with room light, and incubated for additional 8 h in the dark before FACS analysis.

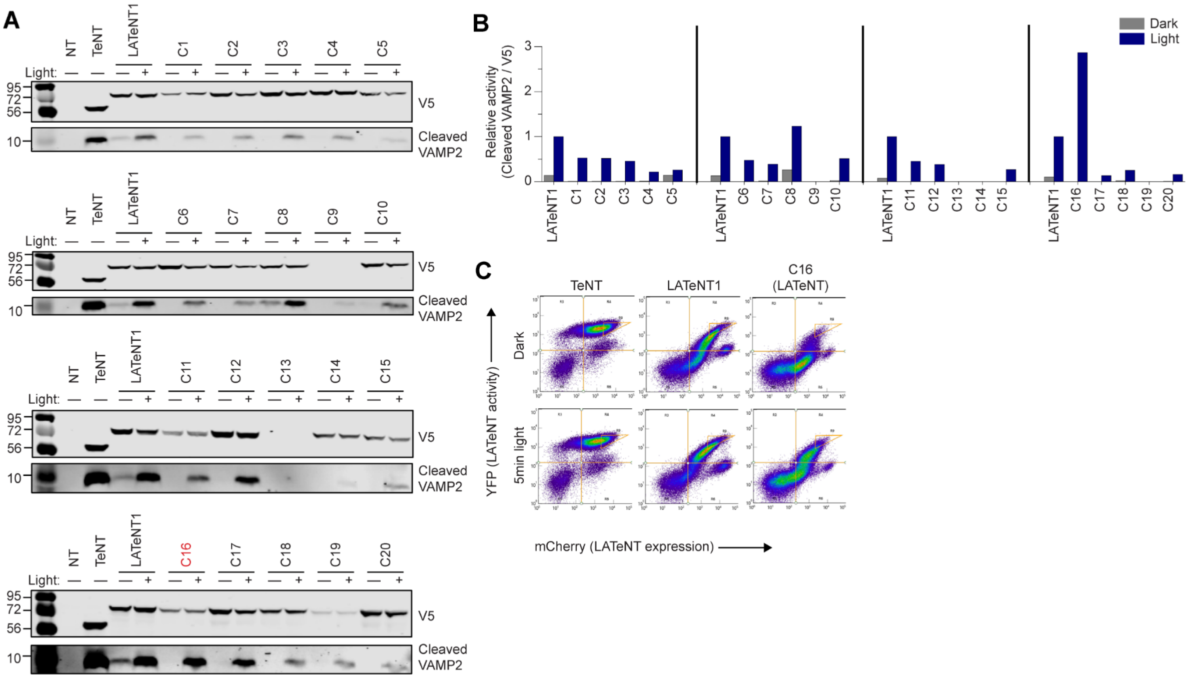


**Supplementary Figure 2. Analysis of clones enriched from directed evolution. Related to Figure 1. (A)** Western blots showing activities of 20 LATeNT mutants isolated from Round 5 library (E1R5). VAMP2 reporter HEK 293T cells expressing each mutant were either kept in the dark or stimulated for 30 min with room light. Multiple blots were quantified in parallel by normalizing against TeNT activity. **(B)** Quantification of dark/light activities of LATeNT mutants in (A). **(C)** Comparison of LATeNT1 and C16 in yeast. Yeast cells expressing LATeNT1 or C16 were kept in the dark or stimulated for 5 min with room light, and incubated for additional 8 h in the dark before FACS analysis. Note that yeast expressing TeNT (positive control) show high YFP expression in both dark and light conditions.

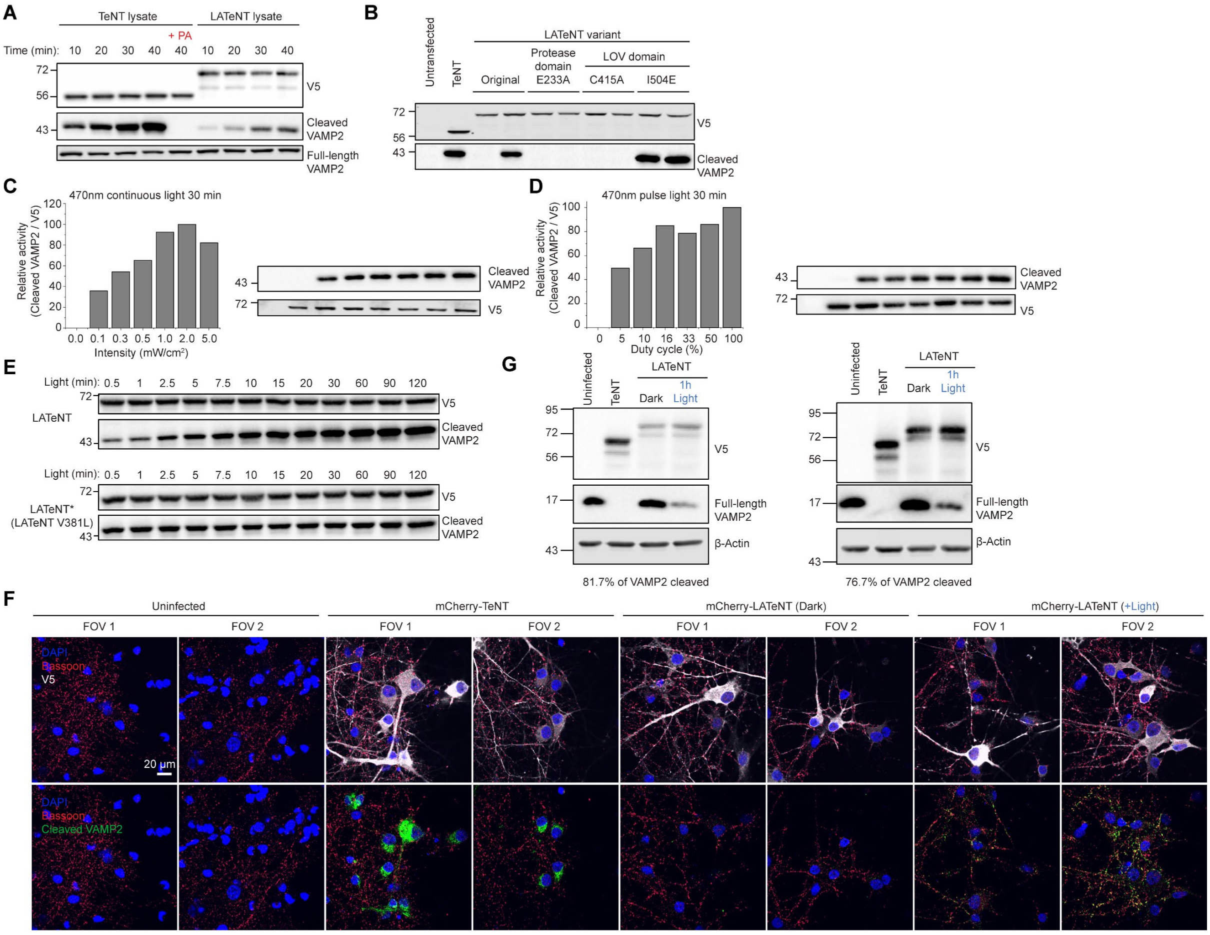


**Supplementary Figure 3. Characterization of LATeNT. Related to Figure 2. (A)** Additional Western blots showing VAMP2 cleavage by TeNT and LATeNT *in vitro*. Adding zinc-chelating picolinic acid completely inhibits TeNT activity (lane 5). Full-length VAMP2 blot shows that the substrate was not limiting. PA, picolinic acid. **(B)** Western blot characterizing effect of point mutations in LATeNT on VAMP2 cleavage in HEK293T cells. Point mutations that inactivate protease activity (E233A)[1], prevent opening of the LOV domain (C415A)[2], or prevent closing of the LOV domain (I504E)[3] were introduced. Light stimulation time was 30 min. This experiment was performed 3 times with similar results. **(C, D)** LATeNT activation with (C) different light intensities or (D) different duty cycles. In (C), VAMP2 reporter HEK293T cells were transfected with LATeNT and stimulated for 30 min with varying intensities of 'continuous' 470 nm blue light. In (D), cells were stimulated for 30 min with 1.0 $mW/cm^2$ 470 nm blue light with different duty cycles (2 s on with 0-38 s off). **(E)** Western blots showing activities LATeNT and LATeNT* with increasing amount of light stimulation. VAMP2 reporter HEK293T cells stably expressing LATeNT and LATeNT* were stimulated for 30 sec to 2 h with 470 nm blue light, and were kept in the dark for the remaining time. **(F)** Additional fields of view showing cleavage of endogenous VAMP2 in cultured neurons. In neurons expressing constitutively-active TeNT, cleaved VAMP2 signal accumulates at the cell body. **(G)** Additional Western blots showing depletion of endogenous VAMP2 in cultured neurons. Neurons expressing LATeNT were stimulated with 470 nm blue light for 1 hour.

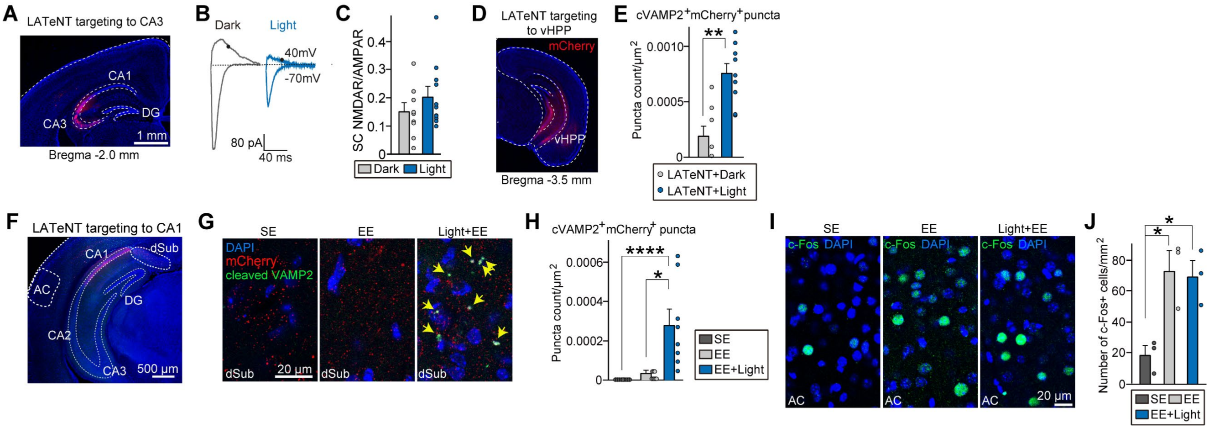


**Supplementary Figure 4. LATeNT inhibits synaptic transmission in the mouse brain, related to Figure 3. (A)** Representative brain section showing mCherry-LATeNT expression in the hippocampal CA3. **(B, C)** Representative traces (B) and summary graphs (C) of NMDAR/AMPAR-EPSC ratios at CA3-CA1 synapses. AMPAR-EPSCs were recorded at -70mV in the presence of picrotoxin, and NMDAR-EPSCs were then recorded at 40 mV ('n' denotes the number of recorded neurons/mice; dark, n = 10/4; light, n = 11/4; Mann-Whitney *U* test). **(D)** Representative brain section showing mCherry-LATeNT expression in the vHPP. **(E)** Summary graphs quantifying cVAMP2⁺mCherry⁺ puncta density in mice expressing AAV-mCherry or LATeNT. Data are presented as mean ± SEM (n = 9 from 3 mice per group; $^{**}p < 0.01$, Mann-Whitney *U* test). **(F)** Representative image showing mCherry-LATeNT expression in the hippocampal CA1. Scale bar = 500 µm. AC, auditory cortex; DG, dentate gyrus; dSub. dorsal subiculum. **(G)** Representative images of dSub regions from mice injected with AAVs expressing mCherry-LATeNT and analyzed via immunofluorescence staining with an anti-cleaved VAMP2 (green) antibody. Yellow arrows mark colocalization of mCherry with cleaved VAMP2 puncta. Scale bar = 20 µm. **(H)** Summary graphs quantifying cVAMP2⁺mCherry⁺ puncta density in mice under SE (dark gray), EE (light gray), or EE+Light (blue) conditions. Data are presented as mean ± SEM (n = 8–9 from 3 mice per group; $^{*}p < 0.05$, $^{****}p < 0.0001$; ANOVA with a non-parametric Kruskal–Wallis test). **(I)** Representative images of c-Fos immunostaining in the AC (control) region. Scale bar = 20 µm. **(J)** Quantitative analysis of data from experiment in (E). Data are presented as mean ± SEM (n = 3 mice each after averaging data from 4 sections/mouse; $^{*}p < 0.05$; ANOVA with Tukey's *post hoc* test).

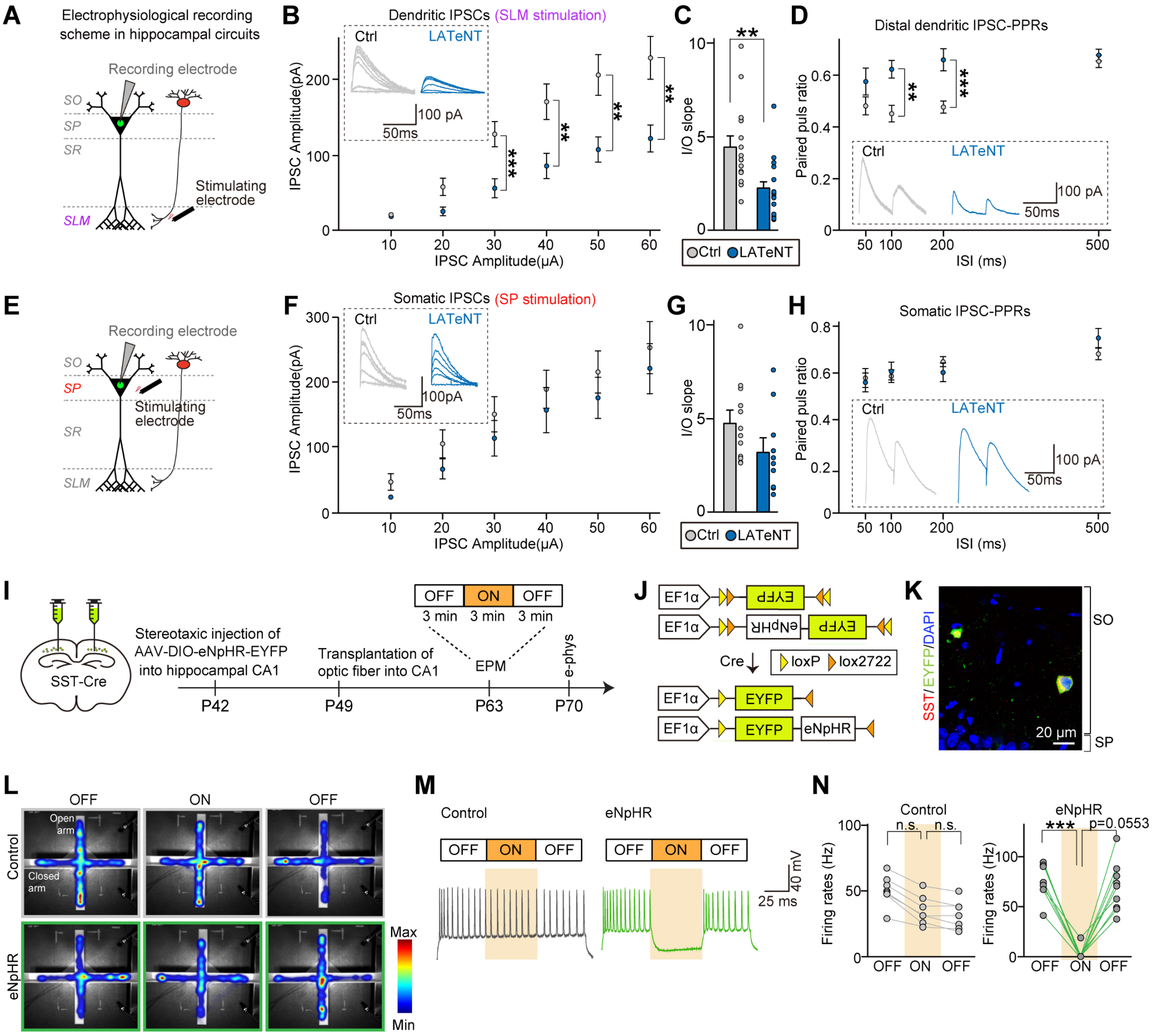

**Supplementary Figure 5. LATeNT reveals causal role for CA1 SST$^{+}$ interneurons in regulating anxiety-like behaviors in mice, related to Figure 4. (A)** Schematic showing recording at CA1 pyramidal neurons with stimulating electrode at SLM layer (dendritic stimulation). SO, stratum oriens; SP, stratum pyramidale; SR, stratum radiatum; SLM, stratum lacunosum-moleculare. **(B, C)** Representative dendritic eIPSC traces, average eIPSC I-O curve (B), and average eIPSC I-O slope (C) from hippocampal CA1 pyramidal neurons ('n' denotes the number of recorded neurons; Ctrl, n = 15; LATeNT, n = 17; **p < 0.01, ***p < 0.001, Mann-Whitney *U* test). **(D)** Representative dendritic PPR traces and average PPR. Data are presented as mean ± SEM (Ctrl, n = 13; LATeNT, n = 15; **p < 0.01, ***p < 0.001, Mann-Whitney *U* test). **(E)** Same as G, with stimulating electrode at SP layer (somatic stimulation). **(F, G)** Representative traces for somatic eIPSCs, average eIPSC I-O curve (F) and average eIPSC I-O slope (G) from hippocampal CA1 pyramidal neurons (Ctrl, n = 12; LATeNT, n = 10; Mann-Whitney *U* test). **(H)** Representative somatic PPR traces and average PPR. Data are presented as mean ± SEM (Ctrl, n = 12; LATeNT, n = 9; Mann-Whitney *U* test). **(I-N)** SST$^{+}$ interneuron manipulation with halorhodopsin instead of LATeNT. (I) Experimental procedure with eNpHR, Three weeks after AAV injection, EPM tests consisting of 5-min epochs with alternating laser manipulation (OFF-ON-OFF) were performed. (J) Cre-dependent recombination of AAV-delivered genes. (K) Representative image showing eNpHR expression in hippocampal CA1 SST interneurons. Scale bar = 50 μm. (L) Heatmaps representing time spent in each arm of the EPM for mice of each group during each epoch (eNpHR group n = 7, EYFP group n = 7). (M, N) Representative traces (M) and summary graphs (N) showing mean firing frequency (percentage) in eNpHR-EYFP-expressing SST+ neurons induced by current injected before and during yellow-light illumination (594 nm, 20 mW/cm$^2$). Data are presented as means ± SEMs (n = 7–9 from 5 mice per group; ***p < 0.001; Friedman test).

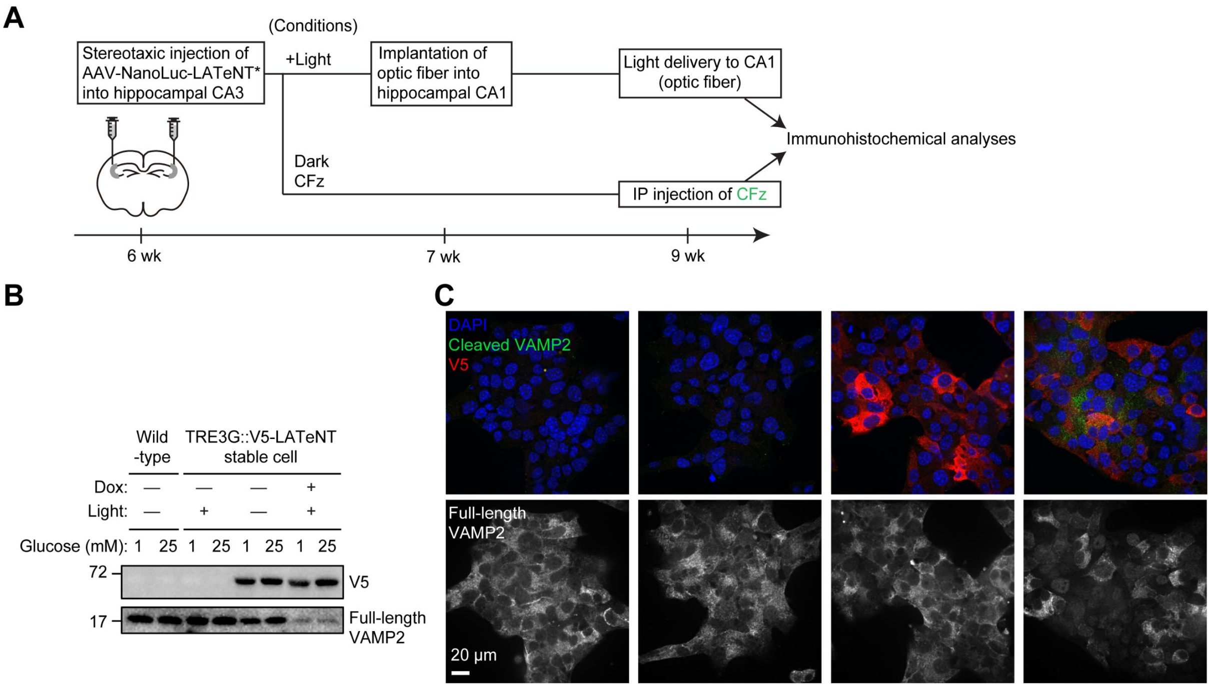


**Supplementary Figure 6. NanoLuc-LATeNT* in vivo and LATeNT for control of insulin secretion.** (A) Experimental procedure for testing NanoLuc-LATeNT* uncaging by cephalofurimazine (CFz) in the mouse brain. AAVs expressing NanoLuc-LATeNT* were injected into the hippocampal CA3 regions. In +light control, optical fibers were implanted into the hippocampal CA1 regions one week after AAV injection, and light was delivered three weeks after AAV injection. For +CFz condition, mice were i.p. injected with 1.3 μmol of CFz three weeks after AAV injection. (B) Western blot for insulin experiment in Figure 5A. (C) Confocal imaging of MIN6 cells from Figure 5A confirms LATeNT expression and light-dependent depletion of endogenous VAMP2.

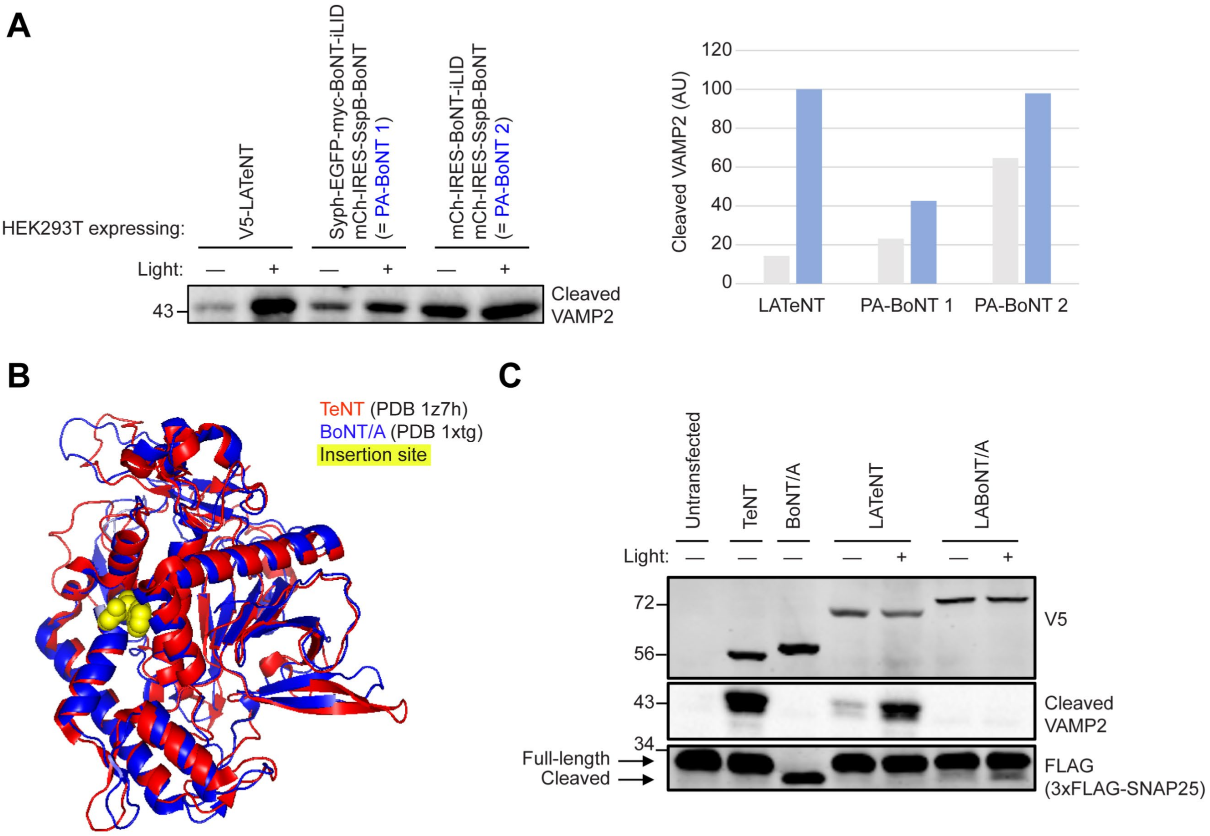


**Supplementary Figure 7. Comparison of LATeNT with PA-BoNT[4] and LOV insertion into BoNT. (A)** HEK293T cells expressing VAMP2 reporter were transfected with LATeNT or PA-BoNT constructs from Liu et al (Addgene plasmids 122981, 122982, and 122985, subcloned under CMV promoter for expression in HEK293T cells). Cells were either maintained in the dark or stimulated for 30 min (for LATeNT) or 4 h (for PA-BoNT) with 14.0 mW/cm$^2$ 470 nm blue light. Left: Western blot analysis of lysates. Right: Quantification of Western blot band intensities. **(B)** Overlaid structures of TeNT protease (PDB 1Z7H) and BoNT/A protease (PDB 3QJ0). LOV domain insertion sites for LATeNT (357/358) and LABoNT/A (347/348) shown in yellow. **(C)** Western blots showing activities of LATeNT and light-activated BoNT/A (LABoNT/A) in HEK293T cells. Cells co-expressed both mCherry-VAMP2 (for LATeNT) and 3xFLAG-SNAP25 (For LABoNT/A). Cells were either maintained in the dark or stimulated for 30 min with 470 nm blue light. Cells expressing constitutively-active TeNT and BoNT/A are included as positive controls. Note that TeNT and LATeNT only cleave VAMP2, while BoNT/A and LABoNT/A only cleave SNAP25.

### Materials

Table of antibodies used in this study.

| Antibody | Source | Vendor | Catalog Number | Dilution(s) |
|---|---|---|---|---|
| Anti-V5 | Mouse | Thermo Fischer Scientific | R96025 | WB: 1:10000; IF: 1:1000 |
| Anti-V5 | Chicken | Nobus Biologicals | NB600-379 | WB: 1:10000 |
| Anti-HA | Rabbit | Cell Signaling Technology | C29F4 | WB: 1:3000; IF: 1:1000 |
| Anti-myc | Chicken | Exalpha Biologicals, Inc. | ACMYC | WB: 1:10000 |
| Anti-VAMP2 | Rabbit | Synaptic Systems | 104 211 | WB: 1:5000 |
| Anti-c-Fos | Rabbit | Cell Signaling Technology | 14609S | IF: 1:500 |
| Anti-Bassoon | Rabbit | Synaptic Systems | 141 118 | IF: 1:1000 |
| Anti-beta actin | Mouse | Santa Cruz Biotechnology | Sc-47778 | WB: 1:3000 |
| Anti-cleaved-VAMP2 (VAMP/B/1148) | Mouse | - | - | WB: 1:5000 |
| Anti-cleaved-VAMP2 (VAMP/B/151) | Mouse | - | - | WB: 1:1000; IF: 1:1000 |
| Anti-GAPDH-HRP | Mouse | Santa Cruz Biotechnology | Sc-47724 HRP | WB: 1:3000 |
| Anti-mouse-AlexaFluor488 | Goat | Invitrogen | A11001 | IF: 1:1000 |
| Anti-chicken-AlexaFluor568 | Goat | Invitrogen | A11041 | IF: 1:1000 |
| Anti-rabbit-AlexaFluor647 | Donkey | Invitrogen | A31573 | IF: 1:1000 |
| Anti-mouse-HRP | Goat | Bio-Rad | 1706516 | WB: 1:5000 |
| Anti-mouse-IRDye 800CW | Goat | LI-COR Biosciences | 92632210 | WB: 1:5000 |
| Anti-mouse-IRDye 680CW | Goat | LI-COR Biosciences | 92668070 | WB: 1:5000 |
| Anti-rabbit-IRDye 800CW | Goat | LI-COR Biosciences | 92632211 | WB: 1:5000 |
| Anti-rabbit-IRDye 680CW | Goat | LI-COR Biosciences | 92668071 | WB: 1:5000 |
| Anti-chicken-IRDye 680CW | Donkey | LI-COR Biosciences | 92668075 | WB: 1:5000 |

### References


1. Li, Y. et al. A single mutation in the recombinant light chain of tetanus toxin abolishes its proteolytic activity and removes the toxicity seen after reconstitution with native heavy chain. *Biochemistry* **33**, 7014-7020 (1994).
2. Swartz, T.E. et al. The photocycle of a flavin-binding domain of the blue light photoreceptor phototropin. *J Biol Chem* **276**, 36493-36500 (2001).
3. Harper, S.M., Christie, J.M. & Gardner, K.H. Disruption of the LOV-Jalpha helix interaction activates phototropin kinase activity. *Biochemistry* **43**, 16184-16192 (2004).
4. Liu, Q. et al. A Photoactivatable Botulinum Neurotoxin for Inducible Control of Neurotransmission. *Neuron* **101**, 863-875.e866 (2019).
